# Cohesin regulates homology search during recombinational DNA repair

**DOI:** 10.1101/2020.12.17.423195

**Authors:** Aurèle Piazza, Hélène Bordelet, Agnès Dumont, Agnès Thierry, Jérôme Savocco, Fabien Girard, Romain Koszul

## Abstract

Homologous recombination (HR) is a ubiquitous DNA double-strand break (DSB) repair mechanism that promotes cell survival. It entails a potentially genome-wide homology search step, carried out along a conserved RecA/Rad51-ssDNA nucleoprotein filament (NPF) assembled on each DSB ends^1–3^. This search is subdued to NPF-dsDNA collision probability, dictated in part by chromatin conformation^2,4^. In contrast to the extensive knowledge about chromatin composition and mobility changes elicited by the DNA damage checkpoint (DDC)^5–7^, whether, how, and to which extent a DSB impacts spatial chromatin organization, and whether this organization in turns influences the homology search process, remains ill-defined^8,9^. Here we characterize two layers of spatial chromatin reorganization following DSB formation in *S. cerevisiae.* While cohesin folds chromosomes into cohesive arrays of 10-20 kb long chromatin loops as cells arrest in G2/M^10,11^, the DSB-flanking regions locally interact in a resection- and 9-1-1 clamp-dependent manner, independently of cohesin and HR proteins. This local structure blocks cohesin progression, constraining the extending NPF at loop base. Functionally this organization promotes side-specific *cis* DSB-dsDNA interactions that scales with loop expansion span, and provides a kinetic advantage for identification of intra- over inter-chromosomal homologies. We propose that cohesins regulate homology search by promoting *cis* dsDNA over-sampling, both upon loop expansion-coupled unidimensional dsDNA scanning, NPF trapping, and chromosome individualization, largely independent of their role in sister chromatid cohesion.

## Main text

How a homologous sequence is identified by a damaged locus in the genome and the nucleus is an enduring conundrum of HR^12^. Here we addressed the spatial dimension of this process, *i.e.* the interplay between chromatin organization and DSB repair by HR using a combination of Hi-C and physical assays for early JM detection in haploid *S. cerevisiae*.

### Cohesins fold chromatin genome-wide following DSB formation

A HO-mediated site-specific DSB was induced on chr. V (**Fig. 1a**). Hi-C libraries were generated at 2 and 4 hours following DSB induction, coincidental with the homology search and DNA strand invasion steps^13^ (**Fig. 1b; Methods**). Hi-C reads coverage and FACS analysis (**Extended Data Fig. 1a-b**) show that 4 hours after DSB induction, ≃2/3^rd^ of the cells are attempting HR repair in a G2/M-arrested stage.

**Figure 1:**
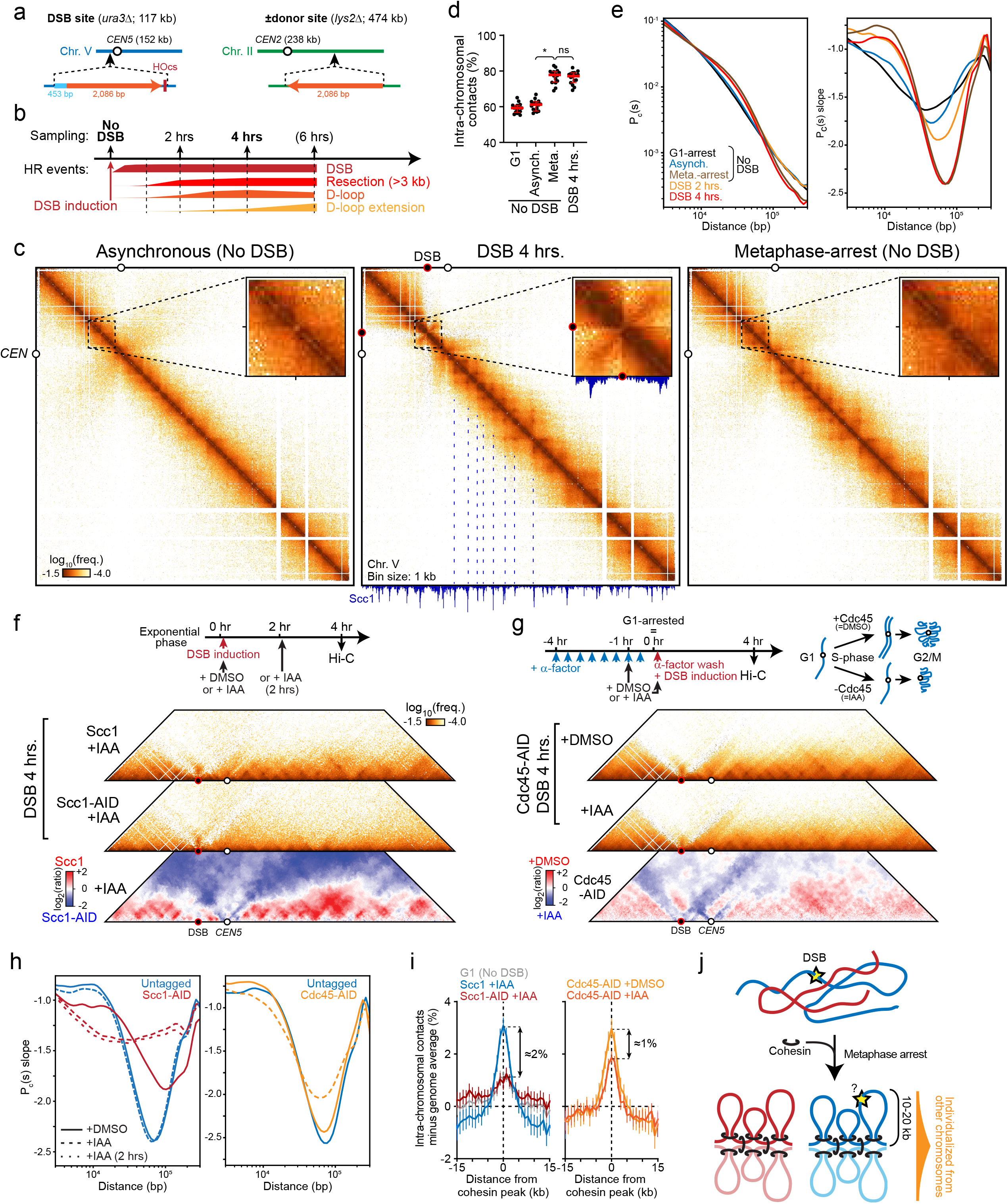
HR repair occurs in a genome spatially reorganized by cohesins as arrays of chromatin loops. **a.** DSB-inducible construct on chr. V, accompanied or not by a region of homology to the left DSB end on chr. II (see below, **Methods**). **b.** Progression of HR as determined in ref. ^5^ following DSB induction in an exponentially dividing cell population and Hi-C sampling times. **c.** Contact maps of chr. V prior to (left) and 4 hours after (middle) DSB induction, or in cells arrested for 4 hours at metaphase without DSB. Magnification of the DSB region is shown. Red dot: position of the DSB. Bin size: 1 kb. **d.** Proportion of intra-chromosomal vs. total contacts and **e.** P_c_(s) (left) and its local derivative (right) in undamaged asynchronous, G1-arrested, and metaphase-arrested cells, or following DSB induction. Data in **d.** represent mean ± interquartile range. **f.** Contact maps 4 hours post-DSB induction in cells with (Scc1+IAA) or without Scc1 (Scc1-AID+IAA). Bottom: Ratio map. **g.** Top: experimental scheme to undergo DSB induction in S/G2 cells lacking a sister chromatid upon depletion of Cdc45 in G1-arrested cells. Middle: contact maps 4 hours post-DSB induction in cells with (Cdc45-AID +DMSO) or without (Cdc45-AID +IAA) sister chromatids. Bottom: ratio map. Representative of biological replicates. **h.** P_c_(s) derivative and **i.** Proportion of intra-chromosomal contacts at and around non-centromeric cohesin peaks (n=474), subtracted from the genome average (dotted line) of data from panels **f**. and **g.** Data in **g.** represent mean ± SEM. **j.** Model for chromosome structure during HR repair. Prolonged metaphase arrest upon DDC activation increases inter-sister loop base promiscuity and connectivity^10^.

Although the typical features of the Rabl chromosome organization were maintained (*i.e.* clustered centromeres and enriched telomere contacts^14^; **Extended Data Fig. 1c**), DSB formation caused drastic reorganization of chromatin at both the local and genome-wide levels (**Fig. 1c-e**, **Extended Data Fig. 1c-e; Methods**). Locally, the DSB region exhibited a local interaction pattern (LIP) consisting in a border centered at the DSB site, and a broad tethering between the left and right DSB regions over ≃25 kb, visible as a enriched contacts perpendicular to the main diagonal and emanating from the DSB site (**Fig. 1c**, see below). Genome-wide, chromatin folded as arrays of loops visible as predominant “triangle-dot” patterns along the main diagonal (**Fig. 1c**), in agreement with the organization of metaphase-arrested cells (**Fig. 1c-e**, **Extended Data Fig. 1b-e; Methods**). Intra-chromosomal contacts increased from 60 to 76%, indicating chromosome individualization (**Fig. 1d**). Contact probability P_c_(s) increased in the 10-60 kb range (**Fig. 1e**), which matched loop length distribution (median: 20 kb, max: 64 kb; **Extended data Fig. 1d**, **Methods**). Finally, pericentromeric regions remodelled as hairpins (**Extended Data Fig. 1e**), indicative of both chromosome biorientation at metaphase^15^ and retention of centromere-kinetochore interactions during HR repair^7^. Both local and genome-wide structuration were not observed upon HO endonuclease expression without a HO cut-site (**Extended Data Fig. 2)**, and were independently confirmed with a different Hi-C protocol, up to 6 hours post-DSB induction (**Extended Data Fig. 3**). A similar reorganization was also observed in diploid cells with a heterozygous, repairable DSB (**Extended Data Fig. 4**).

Loop bases correspond to sites enriched for the cohesin complex (Smc1-Smc3-Scc1) (**Fig. 1c**), as previously observed^10,16^. We first confirmed the dependency of the chromatin loop organization observed 4h after DSB induction on cohesin (**Fig. 1f**). Conditional depletion of the kleisin subunit (Scc1-AID; **Methods**) simultaneously with, or two hours after DSB induction abrogated loop folding and chromosome individualization without affecting DSB processing nor G2/M arrest (**Fig. 1f, h**, and **Extended Data Fig. 5a-d**). Similarly, we verified that preventing replication in the absence of Cdc45, when cells nevertheless reach metaphase^10,17^ (**Fig. 1g**, **Extended Data Fig. 6a-d**, **Methods**), does not prevent loop formation (median: 17 kb) nor chromosome individualization in presence of a DSB (**Fig. 1g**, **Extended Data Fig 6e**). However, the P_c_(s) range, the strength of loop base interactions, and their preferential intra-chromosomal location were reduced (**Fig. 1g-i**, **Extended Data Fig. 6f-g**). These observations suggest that prolonged metaphase arrest leads to more promiscuous inter-sister loop base interactions strongly insulated from other chromosomes (**Fig. 1i**), akin to TAD borders in G2 human cells^18^. Despite cohesin enrichment in the DSB region^19,20^ (**Fig. 1c**), the LIP was retained on a single chromatid and in the absence of cohesin (**Fig. 1f-g**), showing that loop folding and sister cohesion are not involved in DSB end-tethering. In conclusion, HR takes place amidst the structured context of metaphase-like chromosomes organized into 10-20 kb-long chromatin loops following the DDC-mediated metaphase arrest^21^ (**Fig. 1j**).

### Tethering interaction across the DSB region is resection- and 9-1-1 clamp-dependent, and is a cohesin roadblock

The local interaction pattern (LIP) across the DSB region was not specific to its chromosomal context, as it was also observed with a DSB site positioned on chr. IV (**Extended Data Fig. 2**). The pattern extends over time (**Extended Data Fig. 3**) and fits the resection span (**Fig. 1c**, and see below), raising the possibility that this cohesin-independent feature is mediated by repair factors.

We addressed the role of short (Mre11/Rad50/Xrs2 aka MRX complex) and long-range (Exo1 and Sgs1/Dna2) resection factors^22^. The LIP was abrogated in the *mre11Δ, exo1Δ,* and *exo1Δ sgs1Δ* mutants, and in the catalytic-deficient *exo1-D173A* mutant (**Fig. 2a-b**), which exhibited varying degrees of resection initiation or extension defects^18^ (**Extended Data Fig. S7a,c**). Only the *sgs1Δ* single mutant retained the LIP (**Fig. 2b**) and did not exhibit resection defects^23,24^ (**Extended Data Fig. S7a**). Long-range resection both displaces the Ddc1/Mec3/Rad17 clamp (human 9-1-1) away from the DSB end^21,25^ and provides a ssDNA substrate for HR proteins recruitment. Strikingly, *RAD17* or *MEC3* deletion abrogated the LIP without detectable resection defects (**Fig. 2c, Extended Data Fig. 7c-d**), indicating that resection itself does not mediate LIP. The core HR proteins Rad51 and Rad52 were dispensable for LIP (**Fig. 2d**, **Extended Data Fig. 7e-f**). Consequently, we propose that the LIP results from 9-1-1 clamp interactions between the dsDNA at the resection junctions on both DSB ends, with MRX mediating the initial end-tethering^26^ (see below **Fig. 2h**, and **Extended Data 7g**). This interaction may be direct, as Rad17 dimerizes *in vitro*^27^. Note that none of the DSB processing and HR mutants assessed above notably altered genome-wide chromatin folding (**Extended Data Fig. 7h**).

**Figure 2:**
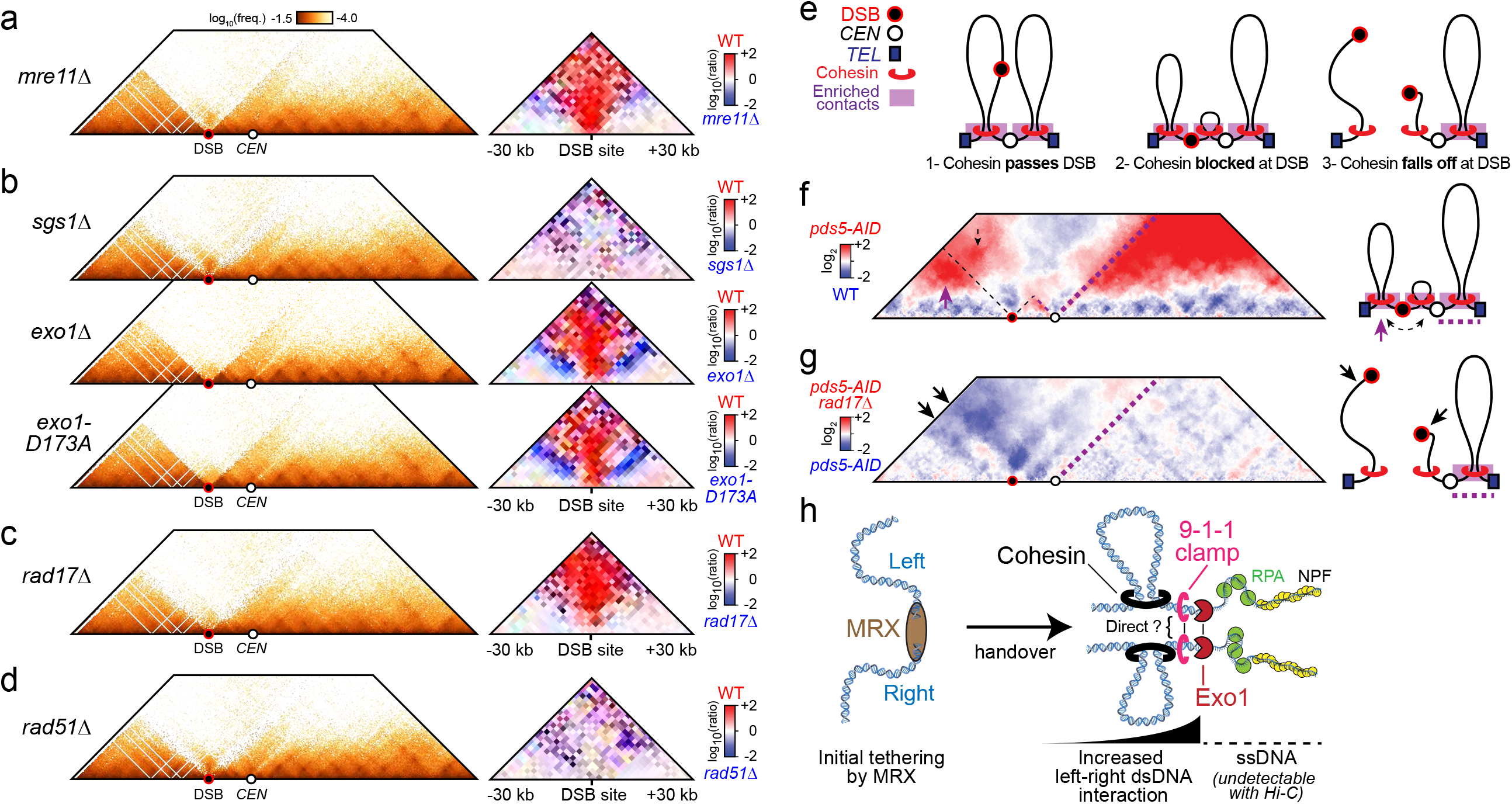
The DSB end-tethering LIP depends on resection factors and the 9-1-1 clamp, and it blocks cohesin progression. **a.-d.** Contact maps (left, bin = 1 kb) and ratio map versus wild-type (bin = 2 kb) of a chr. V region in a *mre11*Δ strain (a), *sgs1*Δ, *exo1*Δ, and *exo1-D173A* strains (b), a *rad17*Δ strain (c), and a *rad51*Δ strain (d). **e.** Possible interaction scenarios in a depending on the cohesin behavior at a DSB. **f.** Ratio map of a *pds5-AID* over a wild-type strain 4 hours post-DSB induction. **g.** Ratio map of a *pds5-AID rad17*Δ over a *pds5-AID* strain 4 hours post-DSB induction. **h.** Model for the DSB end-tethering LIP. Mre11-Rad50-Xrs2 presumably mediates initial end-tethering, subsequently devoted to the Ddc1/Mec3/Rad17 (9-1-1) clamp pushed internally upon resection progression by Exo1^7^. As ssDNA molecules cannot be detected with Hi-C, additional interactions within the resected regions cannot be ruled out.

Finally, we addressed whether the DSB region acts as a cohesin barrier, as proposed in a recent preprint in mammals^28^. To this end we used a hypomorphic mutant (*pds5-AID*) of the cohesin unloader Pds5^29,30^. Pds5-deficient cells feature enlarged loops, which highlights strong cohesin roadblocks, such as centromeres^10^ (**Extended Data Fig. 8a-d**), without DSB processing defects (**Extended Data Fig. 8e** and **9a**). Each of the three possible scenarios (*i.e.* the cohesin can bypass, is blocked, or falls off at the DSB) predicts specific interaction profiles for both the centromere of chr. V and the DSB region (**Fig. 2e**). First, Pds5-deficient cells exhibited an interruption of the contact signal made by the centromere specifically on the left arm on chr. V, ruling out that cohesin can pass the DSB region (**Fig. 2f**). Second, the left DSB end interacted strongly with its distal arm region, consistent with cohesin being blocked at the DSB region rather than falling off (**Fig. 2f, Extended Data Fig. 9b**). This left TEL-DSB interaction was reduced (although not abolished) in the LIP-deficient *pds5-AID rad17Δ* mutant, to the profit of *trans* interactions exceeding that of any single mutant, specifically on this chromosome arm (**Fig. 2h, Extended Data Fig. 9b-d**). These results suggest that the 9-1-1 clamp, in addition to mediate DSB end-tethering, also participates in blocking cohesin translocation. This organization places the resection front (*i.e.* the base of the NPF) at the *cis*-interacting chromosome axis (**Fig. 2h**).

### Cohesins inhibit genome-wide homology search, primarily independent of the presence of the sister chromatid

How do these two interacting layers of chromatin organization affect NPF-dsDNA collision probability, homology search? We leveraged Hi-C contact data made by the dsDNA flanking the DSB region as a low resolution homology search readout, akin to FROS arrays used previously to cytologically monitor DSB mobility^31,32^ (**Fig. 3a, Methods**). Localized HR- and homology-dependent contacts between the DSB region and an ectopic donor located on chr. II could be detected using this approach, which increased as expected in a D-loop disruption *sgs1Δ* mutant^13,33^ (**Fig. 3b**).

**Figure 3:**
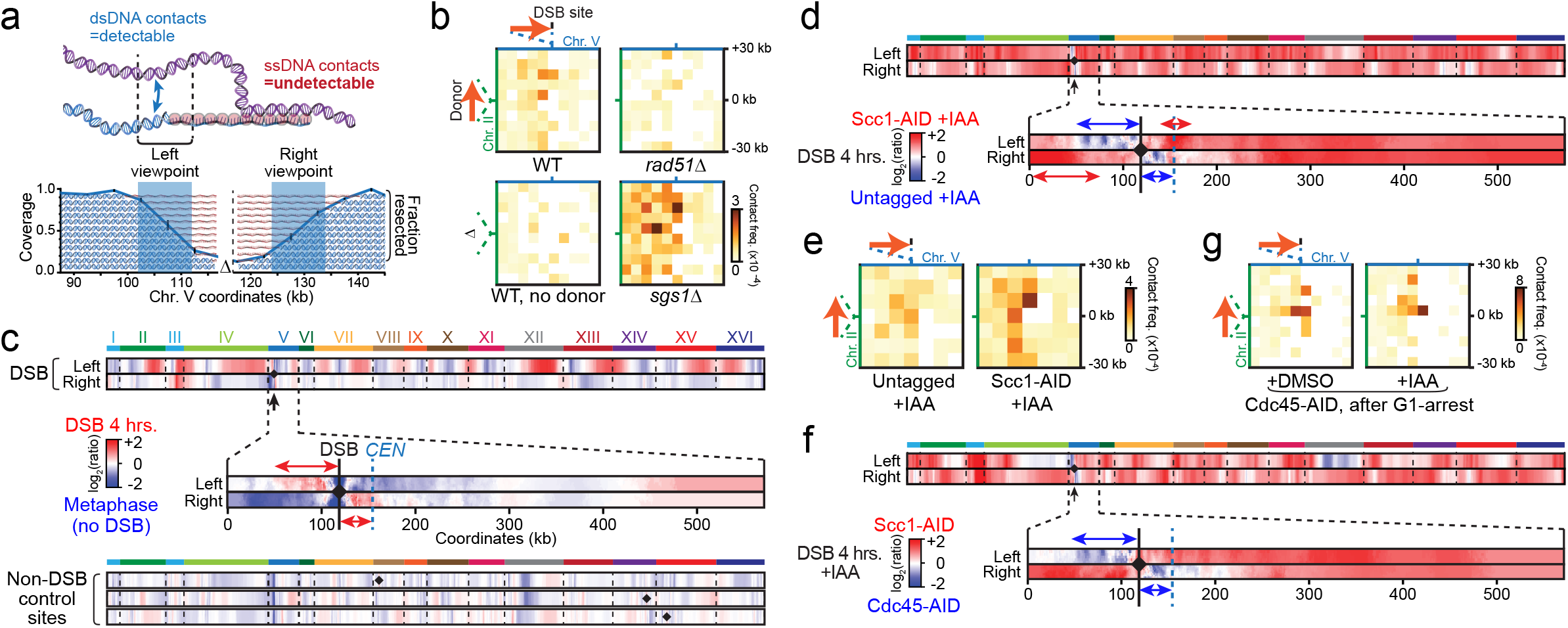
Cohesin inhibits genome-wide homology search, a role mainly independent of the sister chromatid. **a.** DSB-flanking dsDNA contacts are used as homology search for each DSB end (see **Methods** and **Extended Data Fig. 10a**). **b.** Contact maps centered on the DSB on chr. V (x-axis) and a 2 kb-long donor on chr. II (y-axis), unless otherwise stated, in wild-type, *rad51Δ* and *sgs1Δ* strains 4 hours post-DSB induction. Bin size: 6 kb. Linear color scale. **c.** 4C-like ratio plots of the left and right DSB end 4 hours after induction over metaphase-arrested cells (without DSB). The genome-wide (top) and intra-chromosomal (bottom) contact profiles are shown. Black arrow: DSB position. Three undamaged viewpoints, also located at ~35 kb from a centromere, are shown below. Black diamond: location of the viewpoint. **d.-e.** Same as **c.** and **b.**, respectively, but comparing cohesin-depleted (Scc1-AID + IAA) vs. wild-type (Scc1 +IAA) cells. Bin size: 8 kb in **e**. **f.-g.** Same as **c.** and **b.**, respectively, but comparing cells with either two (Cdc45-AID +DMSO) or a single (Cdc45-AID + IAA) chromatid, synchronized after G1-arrest.

A 4C-like representation of dynamically binned contacts made by the left and right DSB viewpoints unveiled their differential interactions across the genome between experimental conditions (**Extended Data Fig. 10a**, **Methods**). Both DSB ends exhibited enriched *cis* contacts in a side-specific manner (60 kb to the left and in the 30 kb DSB-centromere interval on the right) compared to the same undamaged region in metaphase-arrested cells (**Fig 3c**, top). The left DSB region contacted interstitial regions of long chromosomal arms more frequently, at the expense of centromere contacts, a behavior not observed at undamaged control regions nor at the centromere-tethered right DSB end (**Fig. 3c**, bottom). Both ends of a DSB located away from a centromere on chr. IV exhibited increased inter-chromosomal contacts (**Extended Data Fig. 10b**), suggesting that proximity to the centromere anchor inhibits genome-wide homology search. Inter-chromosomal DSB-dsDNA interactions were reduced in HR mutant, but not *cis* interactions (**Extended Data Fig. 10c**). Scc1 depletion, however, radically altered DSB-dsDNA interactions: both DSB ends exhibited a strong and relatively homogeneous increase in contacts with the rest of the genome, at the expense of side-specific *cis* contacts (**Fig. 3e**). DSB-donor contacts also increased 2-fold (**Fig. 3f**). By contrast, the absence of a sister chromatid stimulated only modestly long-range or inter-chromosomal DSB-dsDNA contacts compared to cohesin depletion (**Fig. 3g-h**), accompanied by only a 1.3±0.1-fold increase in DSB-donor contacts (**Fig. 3i**). Cohesin thus promotes intra-chromosomal, side-specific DSB-dsDNA contacts, mainly independent of SCC. This organization inhibits genome-wide DSB-dsDNA contacts and inter-chromosomal donor identification. The NPF only modestly emancipates the DSB region from this constraint.

### Cohesin promotes identification of nearby over inter-chromosomal homologies

The influence of chromatin folding on the kinetics and donor selection process of HR was addressed by quantifying D-loop levels^13^ at ectopic *inter*- or *intra*-chromosomal donors of varying length (**Fig. 4a, Extended Data Fig. 11a**). In the absence of an *intra* donor, *inter* D-loops formed within 3-4 hours following DSB induction before declining, as previously published^13^ (**Fig. 4b**). When present, D-loops formed at the *intra* donor with much faster kinetics and at the expense of *inter* D-loops (**Fig. 4b, Extended Data Fig. 11b**). *Intra* over *inter* donor preference ranged from 15- to 22-fold at equal homology length, and 3- to 4- fold for the 0.6 kb *intra* donor (**Fig. 4c**). Spatial proximity thus confers a strong kinetic advantage for homology identification, only partially overcome by greater homology length.

**Figure 4:**
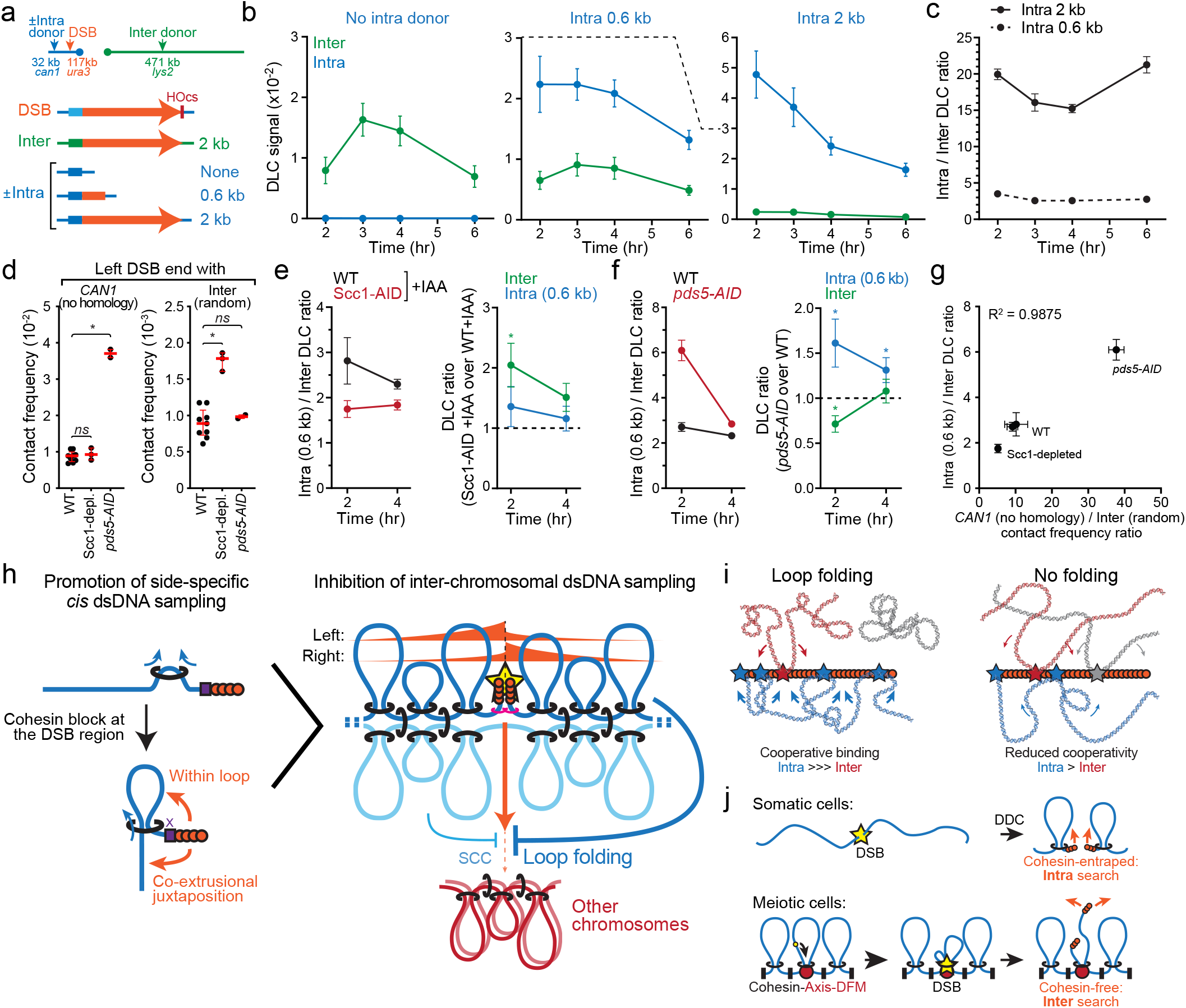
Cohesin promotes homology identification in *cis* and inhibits it in *trans* in multiple ways. **a.** Position of the DSB site, and ectopic intra- and inter-chromosomal donor for competitive D-loop quantification. **b.** Quantification of D-loops formed at the intra- and inter-chromosomal donor sites in strains bearing either no (left), 0.6 kb (middle) and 2 kb (right) of intra-chromosomal homology to the left DSB end. Beware that the y-axis is greater in the right panel than in other panels. Data show mean ±SEM of a biological triplicate. **c.** Ratio of *intra* over *inter* D-loop levels in cells bearing a 0.6 kb- or a 2 kb-long intra-chromosomal competitor. Data show mean ±SEM of a biological triplicate. **d.** Hi-C contact frequency (bin = 10 kb) between the left DSB end and the *can1* locus without a donor (left) or with 100 random inter-chromosomal regions (right) in wild-type, Scc1-depleted and *pds5-AID* cells at 4 hours post-DSB induction. Each point represents a value (left) or a median of values (right) from independent Hi-C experiments. In red: median and interquartile range. **e.** Left: ratio of *intra* over *inter* D-loop levels in cells depleted (Scc1-AID + IAA) or not (WT + IAA) for cohesin. Right: Fold change of *intra* and *inter* D-loop levels upon Scc1 depletion. The *intra* donor is 0.6 kb-long. Data show mean ±SEM of five biological replicates. **f.** Same as in (**e**), comparing wild-type and *pds5-AID* cells. Data show mean ±SEM of three and six biological replicates for wild-type and *pds5-AID* cells, respectively. **g.** *Intra* over *inter* donor preference as a function of Hi-C contact frequency. DLC data are at 2 hours and contact data at 4 hours. Data show mean ±SD. **h.** Model for the three regulations of homology search during DSB repair by HR imparted by cohesins. Right: Cohesin-mediated loop expansion that leads to chromosome folding also promotes side-specific DNA sampling. Right: Chromosome individualization resulting from loop folding favors intra-over inter-chromosomal dsDNA sampling and donor identification. Presence of the sister chromatid (and *a fortiori* SCC) plays a secondary inhibitory role. **i.** Loop folding is expected to cooperatively promote interaction between the NPF and a dominant dsDNA substrate by inter-segmental contact sampling. **j.** Opposite timing of DSB and loop folding in mitosis and meiosis is expected to lead to different DSB-cohesin configurations, which may participate in the opposite HR template preference despite a similar spatial chromatin organization. DFM: DSB-forming machinery.

Conveniently, the contact frequency between the left DSB end and the *intra* donor insertion site (*can1* in the absence of homology) remains unchanged upon cohesin depletion, but increases 4-fold in a *pds5-AID* mutant (**Fig. 4d**, **Extended Data Fig. 11c**). Conversely, the contact frequency between the left DSB end and random *inter* loci increases 2-fold upon loss of chromosome individualization in cohesin-depleted cells, but remains unchanged in a *pds5-AID* mutant (**Fig. 4d**, see also above **Fig. 3d**). This system thus enabled distinguishing two potential roles of cohesin in regulating homology search: the inhibition of *inter* contacts upon Scc1 depletion, and the promotion of *cis* contact upon loop expansion in a *pds5-AID* mutant.

Depletion of Scc1 significantly decreased the *intra* over *inter* donor preference by failing to inhibit *inter* donor identification (**Fig. 4e**, **Extended Data Fig. 11d**), consistent with contact data (**Fig. 3d-e, 4d**). In contrast, *intra* over *inter* donor preference increased in a *pds5-AID* mutant, upon both promotion of *intra* and inhibition of *inter* donor invasion (**Fig. 4f**). This effect was quantitatively more pronounced 2 hours post-DSB induction, coincidental with the establishment of loop folding by cohesin-mediated loop expansion (**Fig. 4f**). Overall, donor preference correlated well with relative contact probability in these three situations (**Fig. 4g**, **Extended Data Fig. 11e**). These results establish that cohesin regulates HR donor identification, both by inhibiting inter-chromosomal contacts and by promoting *cis* contacts. The timing effect observed in the *pds5-AID* mutant (**Fig. 4f**) further suggests that juxtaposition of the NPF with *cis* dsDNA upon loop expansion, and not solely the resulting loop structure, promotes *cis* homology identification (**Fig. 4h** and discussion below).

## Discussion

Here we show that HR repair in *S. cerevisiae* occurs in a chromatin context spatially reorganized at the global and local levels by cohesins and resection-associated factors, respectively. We suggest here plausible ways by which these active organizational mechanisms and their interaction impinge on the homology search process, all contributing to disproportionately favor *cis* dsDNA sampling and homology identification (**Fig. 4h**). First, it localizes and traps the extending NPF at loop base, constraining its explorational capacity to the DSB-borne region, compatible with the inhibitory role cohesin exerts on DSB mobility^7,34^. NPF of increasing length may emancipate from this constraint over time, resulting in a progressive decrease in the preference for *intra* over *inter* donor (**Fig. 4c**). Second, cohesin promotes side-specific NPF-dsDNA interactions and donor identification over the loop expansion tract. It can conceivably result from a juxtaposition of the nascent NPF to dsDNA being extruded at the level of the cohesin ring, or by promoting diffusive contacts within the resulting loop. Third, cohesin-mediated chromosome individualization reduces overall inter-chromosomal contact probability and donor identification. Given the inter-segmental NPF-dsDNA association^2^, this individualization is likely to cooperatively promote *cis* dsDNA sampling (**Fig. 4i**). The sister chromatid, if not competent for repair^35^, acts as a secondary decoy to *trans* NPF-dsDNA contacts. These overlaid constraints inhibit *trans* homology identification by the NPF while promoting it in *cis*, particularly within the loop expansion span (**Fig. 4f**). Cohesin is thus a master regulator of target search during HR, by regulating the NPF-dsDNA contact probability in multiple ways.

Biasing homology search in *cis* is expected to safeguard genome stability against both (i) loss of heterozygosity by inhibiting inter-homolog recombination^36^, and (ii) ectopic repair by occluding most of the repeated content of the genome. If ectopic repair nonetheless occurs, it will be mainly restricted intra-chromosomally, thus mitigating the extent of illegitimate HR repair. The preferential *cis* homology search imparted by cohesin may actually promote formation (and subsequent elimination) of repeat-mediated segmental tandem duplications, which enable rapid adaptation to various environmental stress^37–39^. Such frequent gain and loss events are considered as one of the major evolutionary forces in human^40^. Cohesin *cis* and *trans* regulators, by tuning loop expansion span, may impart an additional layer of control on this opportunistic damage-induced adaptive process.

Interestingly, HR repair in somatic cells parallels the canonical meiotic situation, which also takes place in the context of structured chromosomes^41^ with similar loop patterns, notably during the homology search time frame of zygotene^42,43^. However, meiotic HR exhibits an opposite repair template preference, towards the homologous chromosome rather than the sister chromatid^44,45^. We suspect that these opposite biases may originate from the relative timing of loop establishment and DSB formation^46^, leading to opposite NPF-cohesin configurations and potentially altered NPF dynamics (**Fig. 4i**, and **Supplementary discussion**).

In conclusion, our work reveals the local and genome-wide spatial reorganization of chromatin that accompanies the outcome of a DSB in cycling cells and its functional impact on homology search during HR repair. This minute yet essential DNA metabolic process is subordinated to collision probabilities with other genomic loci, which is partly dictated by cohesins and its interaction with local DSB processing factors, with implications for genome maintenance, evolution and adaptation, and meiosis. It reinforces the idea that cohesin acts as a general facilitator of target search in other processes, both by promoting long-range 1D scanning upon loop expansion^47,48^, or local 3D “hopping” and inter-segmental sampling by reconfiguring the coiling state of chromatin^2,49^.

## Acknowledgments

We thank Axel Cournac, Cyril Matthey-Doret, Vittore Scolari, Jacques Serizay and Théo Foutel-Rodier for their help with bioinformatics analysis and sharing unpublished scripts and programs, Christophe Chapard for his help with cell synchronization, ChIP and test datasets, as well as members of the Koszul lab and of the Parisian Yeast club for stimulating discussions. We are grateful to Nancy Kleckner, Wolf-Dietrich Heyer and David Leach for helpful comments on the manuscript.

This research was supported by the CNRS as part of its Momentum programme to A.P., and the European Research Council (ERC) under the European Union’s Horizon 2020 to A.P. and R.K. (ERC grant agreement 851006 and 771813, respectively).

## Author contributions

Conceptualization: AP and RK; Experiments: AP, HB, AD, FG, AT; Data analysis: AP, HB, JS; Data interpretation: AP, HB and RK; Supervision: AP and RK; Funding Acquisition: AP and RK; Writing – original draft: AP and HB; Writing – final draft: AP, HB, and RK.

