## Supplementary files for "Cohesin regulates homology search during recombinational DNA repair"

#### Supplementary discussion

##### Homology search regulation in mitosis and meiosis: similar chromatin folding but opposite repair templates.

HR repair in somatic cells parallels the canonical meiotic situation, which also takes place in the context of structured chromosomes[1] that exhibit similar loop patterns, particularly during the homology search time frame of zygotene[2,3]. Highlighting the importance of chromatin structure in regulating homology search, homolog bias depends on Rec8, the meiosis-specific Scc1 paralog[13]. However, meiotic HR exhibits an opposite repair template preference, towards the homologous chromosome rather than the sister chromatid[4,5]. We suspect that these opposite biases may originate from the relative timing of loop establishment and DSB formation, which leads to opposite NPF-cohesin configurations (**Fig. 7C**). Indeed, meiotic loop folding occurs prior to, or concurrently with, DSB formation[15]. The DSB-forming machinery (DFM) physically and functionally interacts with chromosome axis proteins whose co-distribution along chromosomes depends on their physical interaction with Rec8[1,6–8]. These cohesin-, axis-, and DFM-enriched sites correspond to DSB cold-spots while the intervening loop region is "hot"[9]. This conundrum posited a subsequent recruitment of loop sites at the DFM-cohesin-axis sites[8,9] leaving the DSB ends free from cohesin entrapment (**Fig. 4i**). This difference will have opposite consequences on homology search: while cohesin holds the NPF and maintains it preferentially at loop bases in mitosis, both ends are free to move away from the axis in meiosis (**Fig. 4i**). These chromatin "tentacles", up to ≈15 kb (≈600 nm) in length, grant nucleus-wide homology search to at least one DSB end, as previously proposed[4,8]. This differential freedom of the two NPFs imposed by their respective loop arm length may account for (i) their different mobility behavior observed cytologically[10], (ii) their differential ability to invade the homolog[11], and (iii) the requirement for synaptonemal complex-mediated homologs tethering and/or loop enlargement by controlled cohesin release to spatially reunite them for downstream crossover resolution[4]. Hence, regulating the initial NPF-interacting partner might alter the chromatin sampling preference and lead to opposite repair template usage, despite a similar spatial chromatin context.

### Online Methods

#### *Saccharomyces cerevisiae* strains

The genotypes of the *Saccharomyces cerevisiae* strains (W303 *RAD5*+ background) used in this study are listed in the **Key Resources Table** and **Table S1**. The *pGAL1-HO* expression system at *trp1* on chr. IV, the mutagenesis of the endogenous HO cut-site at *MAT* on chr. III, the DSB-inducible construct replacing *URA3* on chr. V and the donor sequence replacing *LYS2* on chr. II have been described previously[12]. Briefly, the DSB-inducible construct contains 327 bp of a fragment of the PhiX genome flanked by multiple restriction sites (including *Eco*RI), a 2,086 bp-long sequence corresponding to the first half of the *LYS2* gene (+4 to +2,090), and a 117 bp HO cut-site. It has been introduced at position 116,167 in S288c coordinates, replacing the *URA3* ORF. Alternatively, this DSB-inducible construct has been introduced on chr. IV at position 845,464 in S288c coordinates (**Extended Data Fig. 2**). The intra-chromosomal donor (*aka* competitor) constructs were introduced within *can1-100* on chr. V at position 33,027 in S288c coordinates. The construct contains a *HIS3* gene for selection purposes, either alone ("No competitor") or together with a 578 bp-long (+4 to +582; "0.6 kb competitor") or a 2,086 bp-long (+4 to +2,090; "2 kb competitor") fragment of the *LYS2* gene. An *Eco*RI site is present 233 bp upstream of the donor sequence and used for D-loop Capture (see below).

The methionine-repressible *pMET-CDC20* construct, the construct for expression of the OsTir1 E3-Ubiquitin ligase *his3::pADH1-OsTIR1-9Myc::HIS3*, as well as auxin-dependent degron-containing Scc1(-V5-Pk3-)AID, Cdc45-(FlagX5-)AID and Pds5-(V5-Pk3-)AID constructs have been described previously[13]. The *sgs1::HIS3* deletion was described previously[12]. The *rad51::kanMX*, *rad52::kanMX*, and *exo1::kanMX* deletions were obtained by transformation of PCR products amplified from genomic DNA of the EUROSCARF yeast deletion collection. The *rad57::LEU2* deletion construct was a gift from W.-D. Heyer. The *exo1-D173A* mutation was introduced by transformation of a synthetic gene fragment together with a *TRP1* marker for selection purposes. The *mre11::kanMX* and *rad17::kanMX* deletion were obtained upon short homology ends-out gene targeting of PCR products amplified from Longtine vectors[14]. Gene deletions were verified by PCR and read coverage of Hi-C experiments. Protein tagging were verified by PCR and Western blotting.

#### Culture media

Liquid YPD (1% yeast extract, 2% peptone, 2% dextrose), YEP-lactate (1% yeast extract, 2% peptone, 2% lactate) and Lactate-MET-TRP (0.17% yeast nitrogen base, 0.5% ammonium sulfate, 0.2% synthetic dropout lacking methionine and tryptophan, 2% lactate) were prepared according to standard protocols. Galactose was dissolved with moderate heating and filter-sterilized.

#### Induction of DSB formation in asynchronous cells

Site-specific DSB formation upon galactose-inducible expression of the HO endonuclease has been described in details previously [12,15]. Briefly, an isolated colony picked from a fresh streak on YPD plate was grown overnight with agitation in liquid YPD, and upon saturation diluted in YEP-lactate media. HO expression was induced by addition of 2% galactose to exponentially growing cultures ($OD_{600} \approx 0.5$-1). All cultures were grown at 30°C. In *rad17Δ, pds5-AID rad17Δ,* and *mre11Δ* DDC-deficient mutants and control wild-type or *pds5-AID* cells, nocodazole (10 µg/ml final concentration) was added simultaneously to galactose (2% final concentration) to prevent cellular division upon damage.

#### Induction of DSB formation in cells synchronously released in S-phase

Exponentially growing cultures in YEP-lactate at 30°C were synchronized at the G1/S transition by addition of 1 µg/ml alpha-factor (ProteoGenix, cat. 559627) every 30 minutes for 4 hours. Cells were washed 3 times with 50 mL of pre-warmed YEP-lactate and released in S-phase in YEP-lactate supplemented with 2% galactose to simultaneously induce HO expression. The cell-cycle status was verified by flow cytometry at -4, -1, 0, 2 and 4 hours relative to the S-phase release.

#### Auxin-induced protein depletion

Cdc45-(FlagX5-)AID depletion was induced in cells synchronized with alpha-factor by addition of 2 mM 3-Indole Acetic Acid (IAA, aka Auxin) solubilized in pure DMSO, or the equivalent DMSO concentration (0.35% final) as a control, 1 hour before release from the G1 arrest. Cdc45-depleted cells did not remodel pericentromeric regions as a hairpin due to the loss of bipolar tension[16], a phenotypic expression of the absence of a sister chromatid (**Extended Data Fig. 6d**).

Scc1-(V5-Pk3-)AID depletion was induced in asynchronous cellular culture by addition of 2 mM IAA (or the equivalent DMSO concentration) at the time of DSB induction or 2 hours after.

#### Induction of metaphase arrest in the absence of DNA damage

Metaphase arrest was induced in the same media and for the same duration as DSB-induced cells. Exponentially growing cultures of APY537 (*pMET-CDC20::TRP1*) in lactate-MET-TRP at 30°C were synchronized at the G1/S transition by addition of 1 µg/ml alpha-factor (ProteoGenix, cat. 559627) every 30 minutes for 4 hours. Cells were washed 3 times with 50 mL of pre-warmed YEP-lactate and released in S-phase in YEP-lactate supplemented with 2% galactose and 2 mM methionine. The cell-cycle status was verified by flow cytometry at -4, 0, 2 and 4 hours relative to the S-phase release (data not shown).

#### Flow cytometry

Approximately $10^7$ cells were collected by centrifugation, re-suspended in 70% ethanol and fixed at 4°C for at least 24 hours. Cells were pelleted, washed in 1 mL of 50 mM Sodium citrate pH 7.0, and resuspended in 1 mL 50 mM Sodium citrate pH 7.0. 200 µL of the resuspension was treated with 1 mg/mL RNase A (Euromedex, cat. 9707-C). Following incubation at 50°C for 1 hr or 37°C overnight, cells were pelleted and resuspended in 1.8 mL of 50 mM Sodium citrate pH 7.0 with 2 µM of SYTOX™ Green (Thermo Scientific, S7020), incubated for 1 hr at room temperature in the dark, and sonicated for 10 seconds on a Bioruptor. Flow cytometry profiles were obtained on a MACSQuant machine and analyzed using Flowing Software 2.5.1.

#### Chromatin-immunoprecipitation, sequencing and analysis

Inspired from Hu et al 2015[17]. Fifteen OD600 of *Saccharomyces cerevisiae* cells were mixed with 3 OD600 of *Candida glabrata* exponentially growing cells into a total volume of 45 mL. Each condition is performed in triplicate. Cells were fixed by the addition of 4 mL of the Fixative solution (50 mM Tris-HCl pH 8.0, 100 mM NaCl, 0.5 mM EGTA, 1 mM EDTA, 30% (v/v) formaldehyde) for 30 min at RT and then quenched by the addition of 2 mL of 2.5 M glycine solution for 5 min at RT. After 3 min of centrifugation at 3,500 rpm, cells were washed with ice cold PBS. Cells were resuspended in 300 µL of Chip lysis buffer solution (50 mM HEPES-KOH pH 8.0, 140 mM NaCl, 1 mM EDTA, 1% (v/v) Triton X-100, 0.1% (w/v) sodium deoxycholate, 1 mM PMSF, 1X cOmplete protease inhibitor cocktail (Roche, cat. 11697498001)) and lysed with Precellys in 2 mL tubes (10 times agitation at 7800 rpm during 30 sec - 20 sec stop). 700 µL of Chip lysis buffer solution was added to reach the required volume of 1 mL for sonication by CovarisS220 that shared DNA into 300 bp fragments on average. The soluble fraction was recovered by centrifugation at 13300 rpm at 4°C for 10 min. 80 µL of the supernatant in each condition was saved up and refered as WCE. 5 µg of anti-PK antibody (Abcam, cat. ab27671) was added and incubated in the soluble part ON at 4°C. 50 µL of Protein G dynabeads (Invitrogen, cat. 10004D) were added for 2 hours at 4°C. Beads were then washed 2 times with Chip lysis buffer, then 3 times with Chip lysis buffer higher salt (50 mM HEPES-KOH pH 8.0, 500 mM NaCl, 1 mM EDTA, 1% (v/v) Triton X-100, 0.1% (w/v) sodium deoxycholate, 1 mM PMSF, 1X Complete protease inhibitor cocktail (Roche)), then 2 times with Chip washing buffer (10 mM Tris–HCl pH 8.0, 0.25 M LiCl, 0.5 % NP-40, 0.5 % sodium deoxycholate, 1 mM EDTA, 1mM PMSF) and once with TE buffer (10 mM Tris–HCl pH 8.0, 1mM EDTA, 50 mM NaCl). Elution was performed by incubation of the beads in 120 µL of TES buffer (50 mM Tris–HCl pH 8.0, 10 mM EDTA, 1% SDS) at 65°C for 15 min. Supernatants were collected and referred to as IP. WCE were mixed with 40 µL with TES3 buffer (50 mM Tris–HCl pH 8.0, 10 mM EDTA, 3% SDS). IP and WCE were decrosslinked by overnight incubation at 65°C. RNA was degraded by incubation at 37°C for 1h with 2 µL of RNAse A (10mg/mL, Euromedex, cat. 9707-C). Addition of 10 µL of proteinase K (20mg/mL, Eurobio, cat. GEXPRK01-B5) and incubation at 65°C for 2 hours allowed protein removal. DNA was purified by Phenol:Chloroform:Isoamyl alcohol (Sigma-Aldrich, cat. P3803) extraction followed by isopropanol precipitation, and the pellet resuspended in 60 µL of TE buffer (10 mM Tris-HCl pH 8.0, 1 mM EDTA). Preparation of the samples for paired-end sequencing on an Illumina NextSeq500 (2x35 bp) was performed with TruSeq Nano DNA Library Prep (Illumina, cat. FC-121-9010DOC). Bowtie2 was used for two rounds of alignments, first on *C. glabrata* (CBS138) and then on *S. cerevisiae* allowing the generation of an alignment of IP and WCE that exclusively maped on *S. cerevisiae* (an vice-versa for *C. glabrata*). The obtained SAM file was filtered and converted into a BAM file using Samtools view (parameters -q 5 -F 4). Chip-seq profiles were then normalised by the number of million sequences and multiplied by ORi factor[17] ($WCE_{Cg}IP_{Sc}$ / $WCE_{Sc}IP_{Cg}$, in which $WCE_{Cg}$ and $IP_{Cg}$ correspond to the number of paired reads that mapped uniquely on *C. glabrata* genome and same for *S. cerevisiae* reads) using bedtools genomecov and converted into BigWig using BedToBigWig.

#### Hi-C procedure and sequencing

Cell fixation with 3% formaldehyde (Sigma-Aldrich, Cat. F8775) was performed as described in ref. [13]. Quenching of formaldehyde with 300 mM glycine was performed at room temperature for 20 min. Most Hi-C experiments were performed with a Hi-C kit (Arima Genomics) with a double DpnII + HinfI restriction digestion following manufacturer instructions, except those presented in **Extended Data Fig. 3**, performed as in ref. [13] with a single DpnII restriction digestion. Preparation of the samples for paired-end sequencing on an Illumina NextSeq500 (2x35 bp) or on Illumina Novaseq 6000 (2x150 bp) was performed as described in ref. [13].

#### D-loop Capture assay

D-loop capture assay was performed as in ref. [12], detailed in [15]. The rationale is depicted in **Extended Data Fig 11a**. Briefly, $1.5x10^8$ cells were collected and resuspended in a 0.1 mg/mL Trioxsalen solution (Sigma-Aldrich, Cat. T6137), crosslinked upon 365 nm UV irradiation on a BIO-LINK irradiator (Vilber-Lourmat, Cat. BLX-365) for 10 min, washed and frozen at -20°C. Cells were lysed following spheroplasting with Zymolyase (MP Biomedicals, cat. SKU 08320931) by addition of 0.1% SDS 15 min at 65°C, in presence of 4 pM of APO563 (*i.e.* a long oligonucleotide enabling restoration of a restriction site on the resected broken molecule, see **Extended Data Fig 11a** and **Table S2**). SDS was quenched with 1% Triton X100 and DNA digested in 1X Cutsmart Buffer (NEB, cat. B7204S) with EcoRI-HF (NEB, cat. R3101L) for 60 min at 37°C. The restriction enzyme was heat inactivated 10 min at 55°C in 1% SDS, subsequently quenched with 2% Triton X100. The ligation reaction was performed in dilute condition (≈$1.8x10^4$ genome/µL) with T4 DNA ligase (NEB, cat. M0202L) for 90 min at 16°C. Proteins were digested with 20 µg of Proteinase K (Thermo Scientific, cat. EO0491) for 30 min at 65°C and DNA was purified by Phenol:Chloroform:Isoamyl alcohol (Sigma-Aldrich, cat. P3803) extraction followed by isopropanol precipitation, and the pellet resuspended in 50 µL of TE buffer (10 mM Tris-HCl pH 8.0, 1 mM EDTA) for 30 min at 37°C. Approximately $6x10^5$ genome equivalents (2 µL) are used per qPCR reaction, performed on a Roche LightCycler 96 with the Faststart SYBR Green Master Mix (Roche, cat. 4673484001) according to the manufacturer instructions. Various primer couples were used to control for DNA amount, DSB formation, resection efficiency and digestion/ligation efficiency. Two specific couples enabled determining chimeric products formed between the unique sequences located upstream of the regions of homology at *CAN1* (intra-chromosomal) and *LYS2* (inter-chromosomal). A list of primers used and their purpose is provided in **Table S2**. The DLC values were normalized by the digestion/ligation efficiency and the resected DNA fraction.

#### Protein extraction and western blotting

Protein extracts for western blot were prepared from $5x10^7$ to $10^8$ cells. Cells were lysed in cold NaOH buffer (1.85 N NaOH, 7.5% v/v beta-Mercaptoethanol) for 10 min in ice. Addition of trichloroacetic acid (15% final) for 10 min in ice allowed protein precipitation. After centrifugation 15000 g for 5 min, the pellets were resuspended in 100 µL of SB++ buffer (180 mM Tris-HCl pH6.8, 6.7 M Urea, 4.2% SDS, 80 µM EDTA, 1.5% v/v Beta-mercaptoethanol, 12.5 µM Bromophenol blue). Denaturation was performed by heating 5 min at 65°C. Precleared extracts were resolved on 12% precast polyacrylamide gel (Bio-Rad, cat. 4561043) and blotted on a PVDF membrane (GE Healthcare, cat. 10600023). Membranes were probed with mouse anti-Flag antibody diluted at 1:1000 (Sigma-Aldrich, cat. F3165), or anti-V5 antibody diluted at 1:10000 (Abcam, cat. ab27671), or anti-GAPDH antibody diluted at 1:10000 (Avivasysbio, cat. OAEA00006), and revealed with an HRP-conjugated anti-mouse IgG antibody (Promega, cat. W4028) using the ECL Prime kit (GE Healthcare, cat. RPN2232) and a G:BOX syngene.

#### Processing of the reads, computation of contact matrices, and generation of contact maps

Reads were aligned and the contact data processed using Hicstuff, a custom made pipeline from the lab (https://github.com/koszullab/hicstuff). Briefly, pairs of reads were aligned iteratively and independently using Bowtie2 in its most sensitive mode against the *S. cerevisiae* W303 reference genome (GCA_002163515.1). The reference was deleted for the *URA3* and *LYS2* genes, which are either absent or partially duplicated in our strains, respectively. Each uniquely mapped read was assigned to a restriction fragment. Quantification of pairwise contacts between restriction fragments was performed with default parameters: uncuts, loops and circularization events were filtered as described in ref. [18]. PCR duplicates (defined as paired reads mapping at exactly the same position) were discarded. Contact maps were generated with the "view" function of Hicstuff and normalized using the sequential component normalization procedure (SCN)[18]. Bins were set at either 1 or 2 kb for single chromosomes, or 5 kb for multiple chromosomes, log10 transformed, and rendered.

#### Quantification and comparison of DSB-donor *trans* interactions from Hi-C data and generation of the ratio maps

Two contact matrices to be compared were subsampled to contain the same total number of genome-wide contacts, and binned at 1 kb (dense format). Each row of bins within the 30 kb regions surrounding the DSB site was then subsampled to match the total number of contacts made by the least covered row between the two matrices. Starting from 20 million genome-wide contacts, it represents approximately 60,000 - 100,000 contacts made by the DSB region. Contacts made by the 20 kb on the left side of the DSB with the 20 kb region on each side of the donor site were quantified and compared across samples. Alternatively, for the decompacted Scc1 depletion experiment, 30 kb regions for both the left flanking region of the DSB and each side of the donor site were considered. Matrices were rendered without additional normalization or transformation, binned at either 6 or 8 kb.

#### Generation of ratio maps

Ratio maps in **Fig. 2a-d** and **Extended Data Fig. S7c, e, g** were generated from sparse matrices using the Hicstuff "view" function, normalized, log-transformed and binned at either 1 or 2 kb. Other ratio maps were generated with Serpentine[19]. Briefly, genome-wide contact matrices binned at a 1 kb resolution (dense format), and containing the same number of contacts (upon subsampling of the most covered dataset), were generated with Hicstuff. For analysis of intra-chromosomal interactions, only chromosomes of interest were loaded into Serpentine. Comparison between pairs of contact maps was performed using the default threshold parameters (50 and 5) and the detrending constant determined automatically by Serpentine for each comparison.

For differential analysis of the intra-chromosomal and genome-wide contacts made by 10 kb-wide DSB-flanking regions on chr. V in **Fig. 3 and Extended Data Fig. 10**, each of the 10 rows were subsampled to yield an equal number of contacts per row between each matrix, which allowed to set the detrending constant to 0. This per-row subsampling was used to homogenize data with varying coverages due to resection of the DSB region. In some instances in which overall chromatin folding slightly differed because of inefficient arrest at G2/M, the average default detrending constant determined at undamaged regions was set as baseline in determining left and right DSB contact profiles.

#### Computation of the contact probability as a function of genomic distance

Computation of the contact probability as a function of genomic distance $P_c(s)$ and its derivative have been determined using the "distance law" function of Hicstuff with default parameters, averaging the contact data of entire chromosome arms (see ref. [2] for details).

#### Quantification of coverage from Hi-C data

Quantification of coverage from Hi-C data was determined with custom scripts from sparse contact matrices generated with the Hicstuff "*pipeline*" function. Briefly, the sum of total contact frequencies within 5 kb bins was computed along the broken chromosome for each sample, and normalized onto the average coverage of undamaged cells obtained from a biological replicate.

#### Loop detection and scoring with Chromosight

Chromosight 1.3.1[20] was used to call loops *de novo* from contact maps binned at 1 kb and balanced with Cooler[21]. Matrices were subsampled to contain the same total number of contacts. *De novo* loop calling was computed using the "*detect*" mode of Chromosight, with minimum loop length set at 10 kb and pearson correlation threshold set at 0.4.

#### Generation of aggregated contact maps

Cohesin pile-up contact maps of averaged 29 kb windows were centered on pairs of cohesin-enriched sites and normalized over the average of ten randomly chosen pairs of positions maintaining the same inter-distance range were generated with Chromosight. Cohesin enrichment sites (bin: 1 kb) were determined from Scc1 ChIP-seq profile data from ref.[17] (accession SRR2065069 and SRR2065070). Each bin presenting a >2.5-fold Scc1 enrichment over input was considered as a cohesin enriched site. To avoid biases introduced by centromere regions, sites located within 10 kb from centromeres were discarded, resulting in 474 cohesin-enriched sites.

Centromere pile-up contact maps are the result of averaged 29 kb windows generated by Chromosight centered on centromere position, and normalized over the average of ten randomly chosen pairs of positions along the diagonal.

#### Statistical analysis

The proportion of intra-chromosomal contacts and DLC values were compared using a non-parametric Mann-Whitney Wilcoxon rank sum test, two-tailed. Statistical cut-off was set at 0.05 All statistical tests were performed under R 3.6.2.

#### Data and software accessibility

The accession number for the sequencing reads reported in this study is SRA: SUB7780899.
Open-access versions of the programs and pipeline used (Hicstuff, Chromosight, and Serpentine) are available online on the github account of the lab (https://github.com/koszullab/).
R x64 3.6.1 is available online at https://cloud.r-project.org/.

Bowtie2 2.3.4.1 is available online at http://bowtie-bio.sourceforge.net/bowtie2/.

### Supplementary figure legends

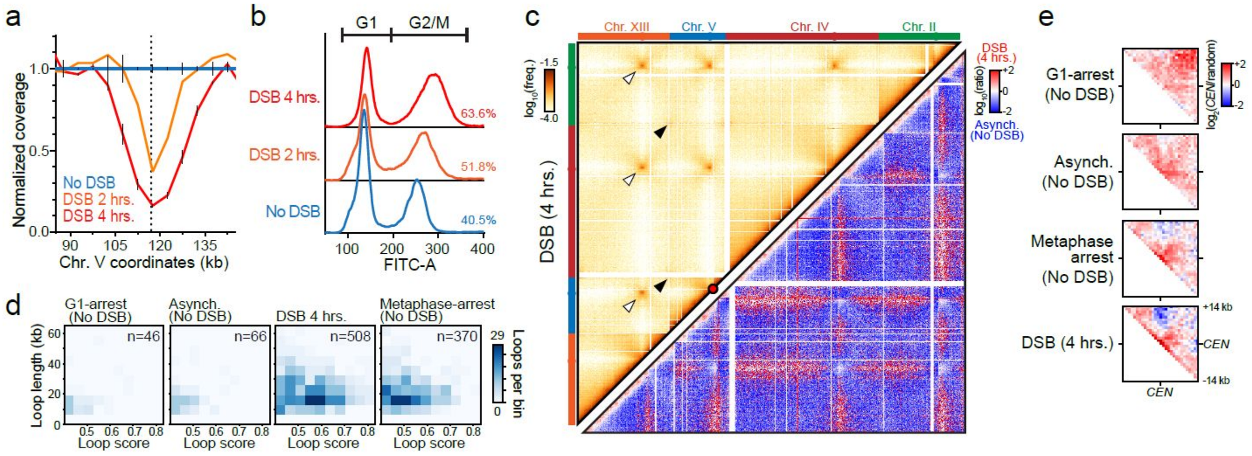


**Figure 1: Genome-wide spatial chromatin reorganization following DSB formation.**

**a.** Hi-C coverage binned at 5 kb prior to and following DSB induction, normalized onto the average coverage in the absence of DSB. Data represent the mean ±SD of a biological replicate, except for DSB 2 hrs (n=1).

**b.** Cell-cycle progression prior to and following DSB induction determined by flow cytometry. The percentage of cells with a 2n DNA content is indicated for each time point.

**c.** Top left corner: Contact map showing intra- and inter-chromosomal contacts made by four chromosomes (II, IV, V, XIII) in a wild-type strain 4 hours post-DSB induction. Examples of inter-chromosomal contacts at the level of centromeres and telomeres are highlighted with white and black arrows, respectively. Bottom right corner: Ratio of contact maps obtained in a wild-type strain 4 hours post-DSB induction (red) over asynchronous cells without a DSB (blue). Bin size: 5 kb for both matrices.

**d.** Heatmap of loop lengths and scores detected with Chromosight in wild-type cells either asynchronous, G1-arrested, or metaphase-arrested without a DSB, or at 4 hours post-DSB induction. The total number of detected loops "n" is indicated. Determined from datasets subsampled at 24 million contacts each.

**e.** Aggregated ratio maps of 29 kb windows centered on centromeres over randomly chosen positions on the diagonal of the contact maps, determined from data presented in **Fig. 1c**. An undamaged G1 control is shown for comparison. Bin size: 1 kb.

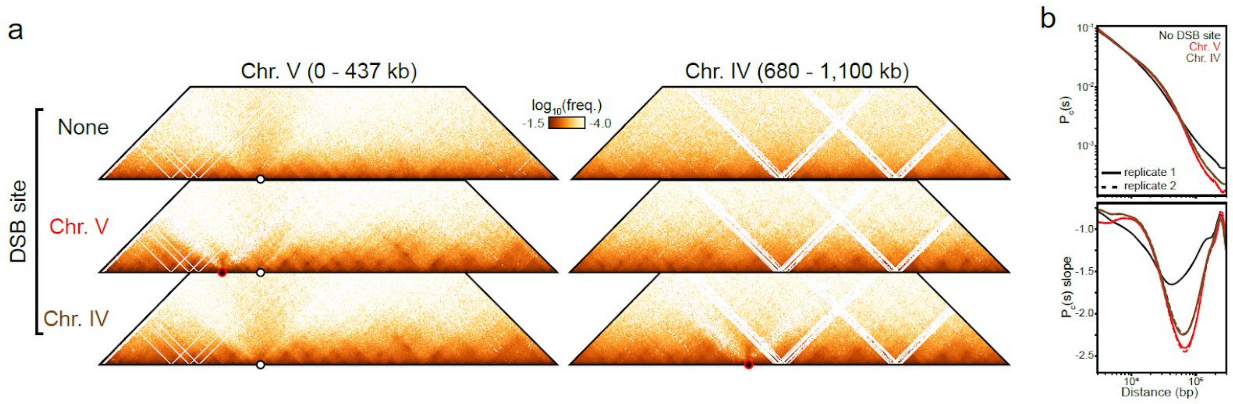


**Figure 2: Local and global spatial chromatin reorganization upon HO expression require a HOcs, but occur irrespectively of its genomic location.**

**a.** Contact maps of regions of Chr. V and Chr. IV in strains bearing either no HOcs (n=1), Chr. V-HOcs (n=2) or Chr. IV-HOcs (n=2) 4 hours post-induction of HO endonuclease expression. The LIP is specific from the DSB position (red dot). Generated from 24 million contacts each. Bin size: 1 kb.

**b.** Contact probability as a function of genomic distance ($P_c(s)$, left) and its derivative (right) determined from samples in (**a**).

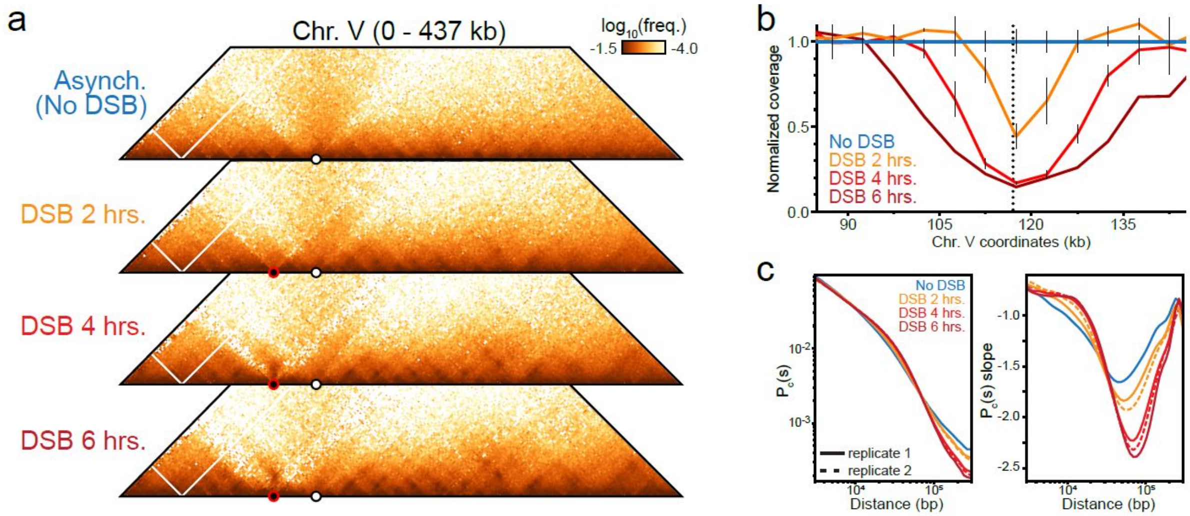


**Figure 3: Local and global spatial chromatin reorganization determined over 6 hours following DSB induction (Hi-C protocol with single DpnII digestion).**

- **a.** Contact maps of a region of Chr. V before and at 2, 4, and 6 hours post-DSB induction (n=3, 3, 4, and 1 biological replicates, respectively). Generated from 17-19 million contacts each. Red dot: DSB position. White dot: centromere. Bin size: 2 kb.
- **b.** Hi-C coverage binned at 5 kb in samples in (a) as in **Extended Data Fig. 1a**. Data represent mean ±SD.
- **c.** Contact probability as a function of genomic distance ($P_c(s)$, left) and its derivative (right) determined from replicates of samples in (a).

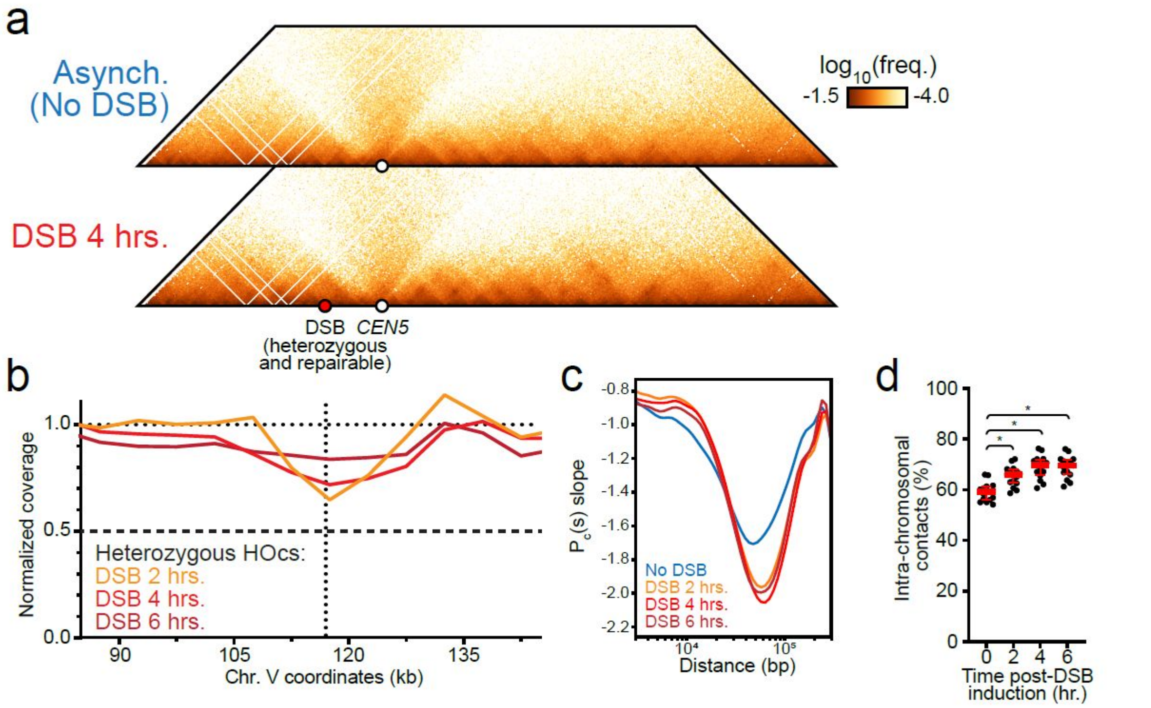


**Figure 4: Spatial genome reorganization upon formation of a repairable DSB in diploid cells.**

- **a.** Contact maps of a region of Chr. V in diploid wild-type cells prior and 4 hours post-induction of a heterozygous DSB (Red dot). Generated from ≃17 million contacts each. White dot: centromere. Bin size: 1 kb.
- **b.** Hi-C coverage binned at 5 kb before and at 2, 4, and 6 hours post-DSB induction as in **Extended Data Fig. 1a**. Coverage drops at 2 hours and is progressively restored at 4 and 6 hours, as inter-homolog repair occurs[15].
- **c.** Derivative of the contact probability as a function of genomic distance $P_c(s)$ before and at 2, 4, and 6 hours post-DSB induction.
- **d.** Proportion of intra-chromosomal vs. total contacts for each 16 chromosomes in diploid wild-type cells before and at 2, 4, and 6 hours post-DSB induction. Red bar: mean ± interquartile range.

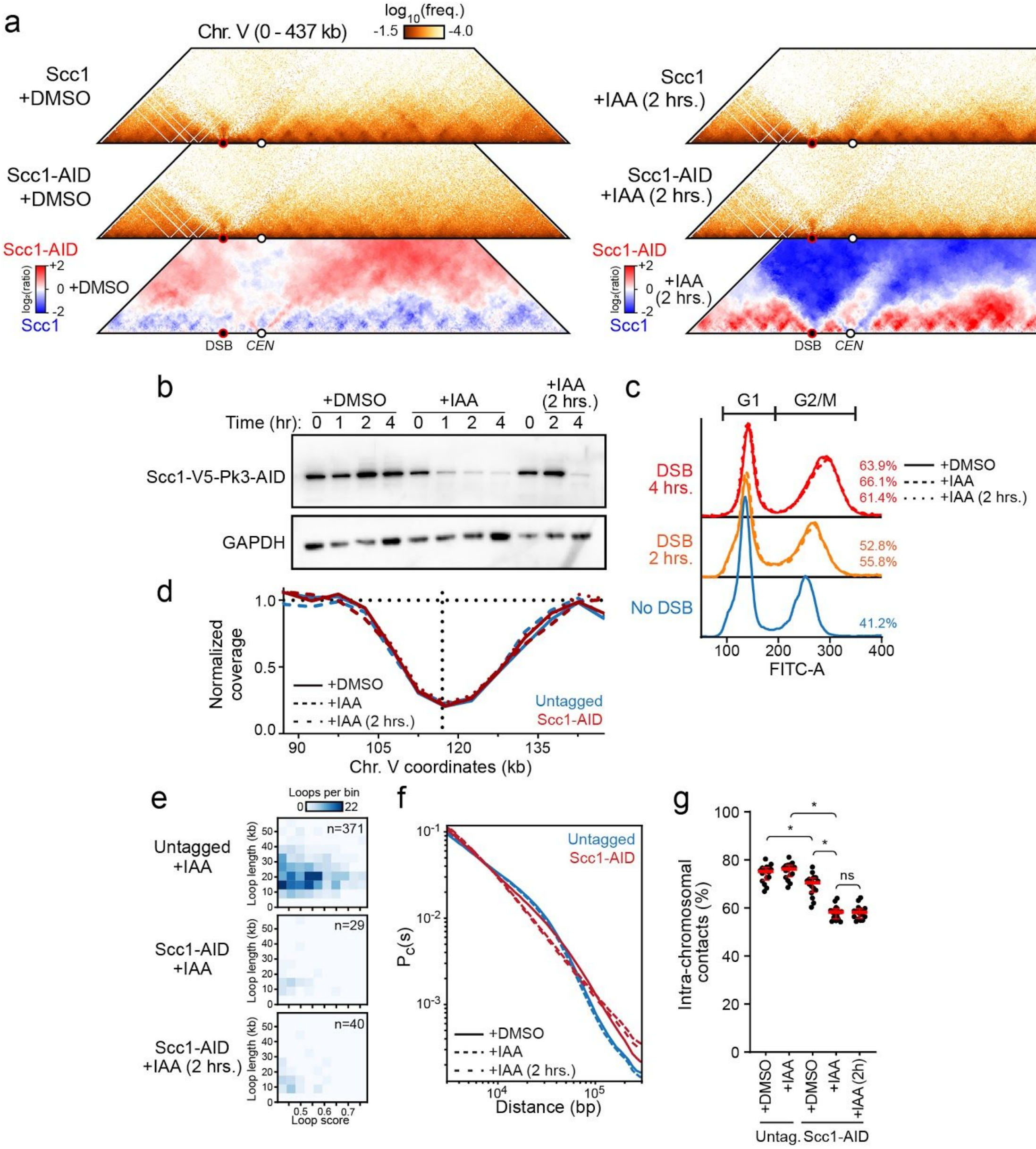


**Figure 5: Loop formation and chromosome individualization following DSB formation is Cohesin-dependent.**

a. Left panels: Contact maps of a region of Chr. V in untagged or Scc1-AID tagged cells at 4 hours post-DSB induction, treated with DMSO added at the same time as DSB induction. Bin size: 1 kb. Bottom left panel: Ratio map of untagged over Scc1-AID tagged cells treated with DMSO at 4 hours post DSB induction. The AID tag mildly alters cohesin regulation, causing loops to become longer. Blue indicates a stronger signal in untagged cells, and vice-versa for red. Right panels: Contact maps of a region of Chr. V in untagged or Scc1-AID tagged cells 4 hours post-DSB induction, treated with IAA added 2 hours after DSB induction, when spatial chromatin reorganization has initiated (**Fig. 1e** and **Extended Data Fig. 3**). Bin size: 1 kb. Bottom right panel: Ratio map of untagged over Scc1-AID tagged cells treated with IAA (2hrs.).

b. Western blot of Scc1-(V5-Pk3-)AID probed with an anti-V5 antibody (top) and a loading control GAPDH (bottom) over the course of IAA or DMSO (mock) treatment of samples presented in (**a**) and **Fig. 1f, h-i**.

c. Cell-cycle progression determined by flow cytometry in samples presented in (**a**) and **Fig. 1f, h-i**. The percentage of cells with a 2n DNA content is indicated for each time point.

d. Hi-C coverage binned at 5 kb in samples presented in (**a**) and **Fig. 1f, h-i**, as in **Extended Data Fig. 1a**.

e. Heatmap of loop lengths and scores detected with Chromosight 4 hours post-DSB induction in control cells (untagged) or in Scc1-depleted cells simultaneously or 2 hours after DSB induction. Determined from datasets subsampled at 15 million contacts each.

f. Contact probability as a function of genomic distance $P_c(s)$ of cells 4 hours post-DSB induction in presence of Scc1 (*i.e.* untagged +DMSO or +IAA) or upon Scc1 depletion (*i.e.* AID tagged, +IAA added simultaneously or 2 hours after DSB induction).

g. Proportion of intra-chromosomal interactions for each 16 chromosomes from samples presented in (**a**) and **Fig. 1f, h-i**. In red: mean ± interquartile range.

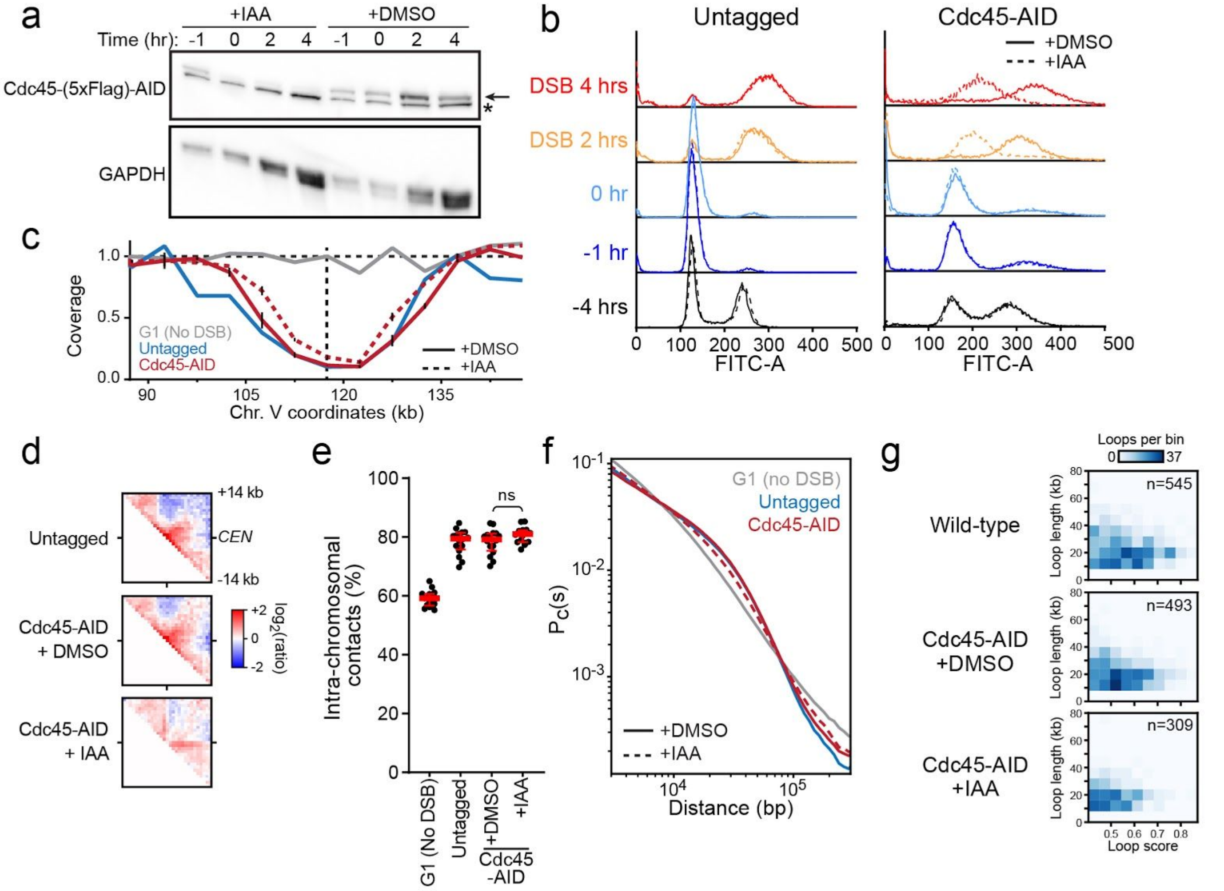


**Figure 6: A sister chromatid is not required for loop formation and chromosome individualization following DSB formation, but contributes to loop base interactions.**

a. Western blot of Cdc45-(5xFlag)-AID probed with an anti-Flag antibody (top) and the loading control GAPDH (bottom) from samples presented in **Figs. 1g-i**. The arrow denotes Cdc45 and the * an aspecific cross-detection by the Flag antibody.
b. Cell-cycle progression determined by flow cytometry in samples presented in **Figs. 1g-i**.
c. Hi-C coverage binned at 5 kb of samples presented in **Figs. 1g-i**, as in **Extended Data Fig. 1a**. Data represent the mean ±SD of a biological replicate.
d. Aggregated ratio maps of 29 kb windows centered on centromeres over randomly chosen positions on the diagonal of the contact maps determined from data presented in **Figs. 1g-i** and from an untagged control.
e. Proportion of intra-chromosomal interactions for each 16 chromosomes from samples presented in **Figs. 1g-i**. In red: mean ± interquartile range. A damaged untagged control and an undamaged G1 control are shown for comparison.
f. Contact probability as a function of genomic distance $P_c(s)$ in DMSO- or IAA-treated Cdc45-AID-tagged or untagged strains 4 hours post-DSB induction. An undamaged G1 control is shown for comparison.

g. Heatmap of loop lengths and scores determined with Chromosight 4 hours post-DSB induction in in Cdc45-AID tagged (+DMSO) and depleted (+IAA) cells, as well as in untagged cells shown for comparison. Loop lengths and scores decrease significantly in the absence of the sister chromatid. Determined from datasets subsampled at 24 million contacts each.

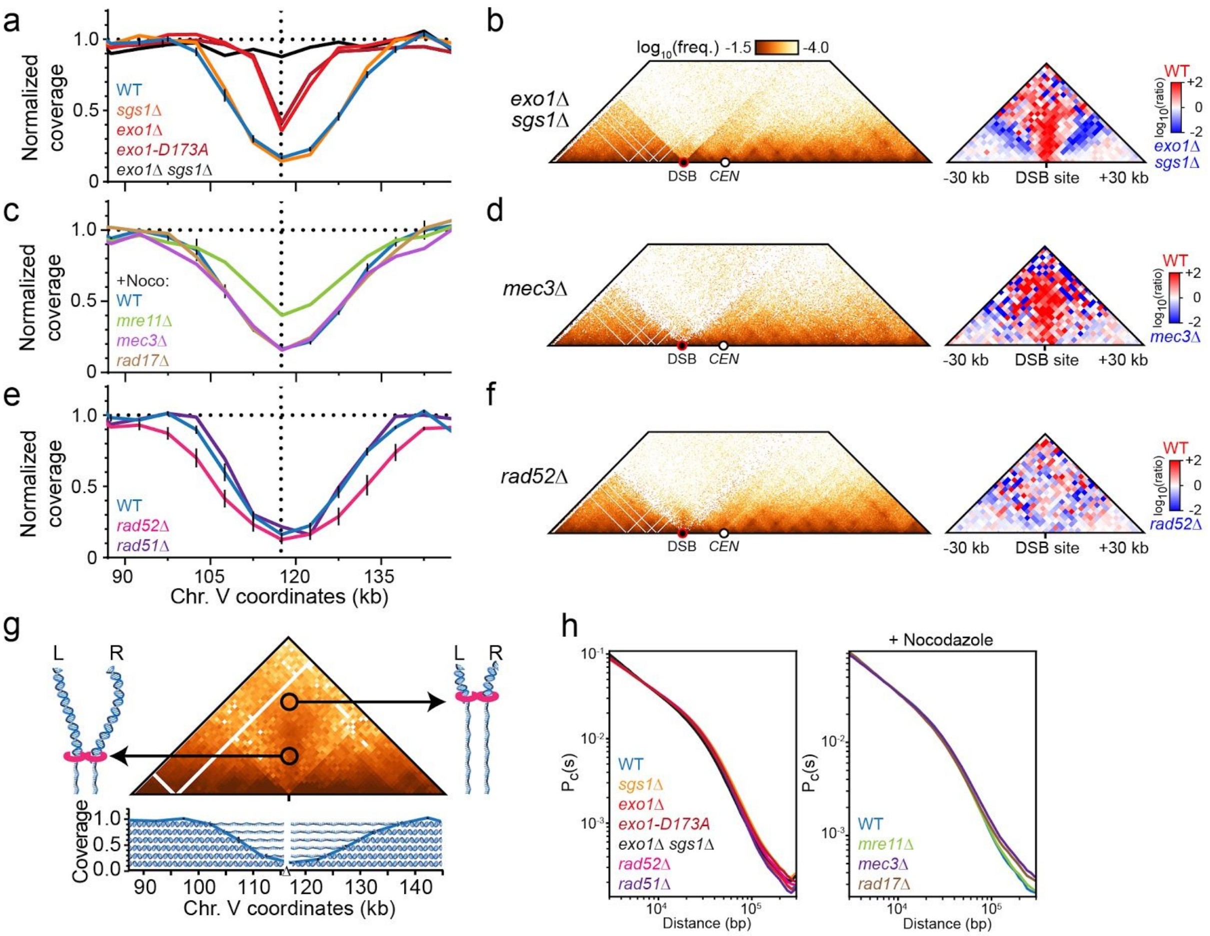


### Figure 7: Determinants and organization of the LIP.

- **a.** Hi-C coverage binned at 5 kb in a wild-type strain and resection mutants. Data are mean ±SEM when available.
- **b.** Contact maps (left, bin = 1 kb) and ratio map versus wild-type (bin = 2 kb) of a chr. V region in a *exo1Δ sgs1Δ* strain.
- **c.** Hi-C coverage binned at 5 kb in nocodazole-treated wild-type, *mre11Δ*, *mec3Δ*, and *rad17Δ* strains. Data are mean ±SEM when available.
- **d.** Contact maps (left, bin = 1 kb) and ratio map versus wild-type (bin = 2 kb) of a chr. V region in a *mec3Δ* strain.
- **e.** Hi-C coverage binned at 5 kb in a wild-type strain and HR mutants. Data are mean ±SEM when available.
- **f.** Contact maps (left, bin = 1 kb) and ratio map versus wild-type (bin = 2 kb) of a chr. V region in a *rad52*Δ strain.
- **g.** Model accounting for the distribution of contacts between the left and right regions surrounding the DSB site. The extent of the contact signal corresponds to the extent of resection. Contact map and coverage at 4 hours post-DSB induction as in **Fig. 1c** and **Extended Data Fig. 1a**, respectively.

**h.** Contact probability as a function of genomic distance $P_c(s)$ in untreated wild-type, *sgs1Δ*, *exo1Δ*, *exo1-D173A*, *exo1Δ sgs1Δ, rad51Δ*, and *rad52Δ* strains (left), or in nocodazole-treated wild-type, *mre11Δ*, *mec3Δ*, and *rad17Δ* strains (right).

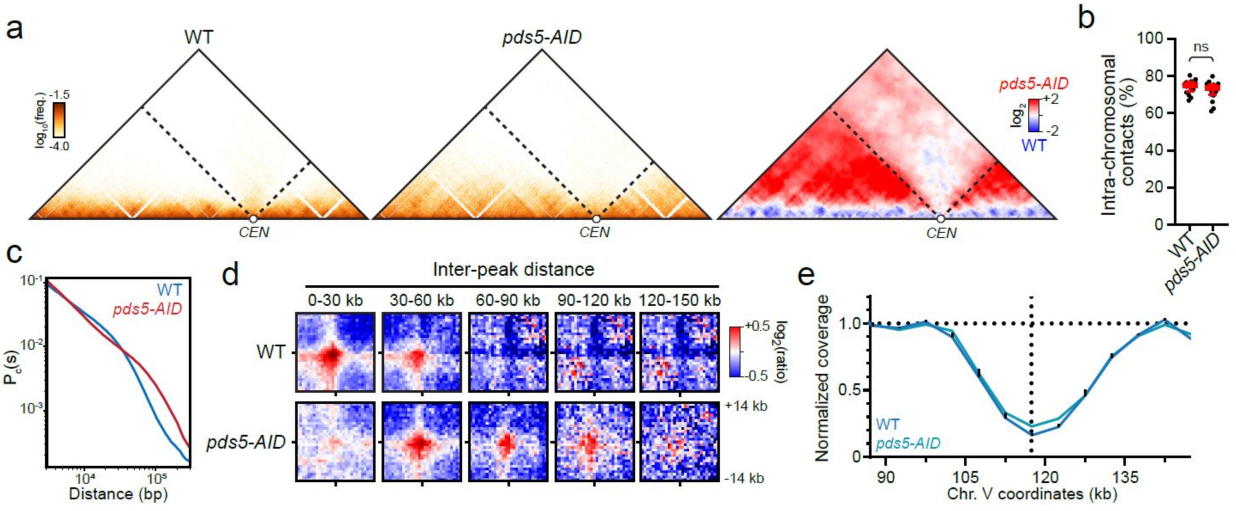


**Figure 8: Loop length increases in a *pds5-AID* mutant, highlighting strong cohesin roadblocks.**

- **a.** Contact maps (bin size: 1 kb) of a region of the undamaged Chr. XI in wild-type and hypomorphic pds5-AID strains 4 hours post-DSB induction on chr. V. Generated from datasets containing ≃15-16 million genome-wide pairs each. Right panel: ratio map of the pds5-AID mutant over a wild-type strain. Dotted lines emanating from *CEN11* highlight intra-chromosomal contacts made by the centromere.
- **b.** Proportion of intra-chromosomal interactions for each 16 chromosomes in wild-type or *pds5-AID* strains 4 hours post-DSB induction. In red: mean ± interquartile range.
- **c.** Contact probability as a function of genomic distance ($P_c(s)$, left) and its derivative (right).
- **d.** Aggregated ratio maps of 29 kb windows centered on pairs of cohesin-enriched over randomly chosen positions in the same inter-distance range. Bin size: 1 kb.
- **e.** Hi-C coverage binned at 5 kb. Data are mean ±SD, when available.

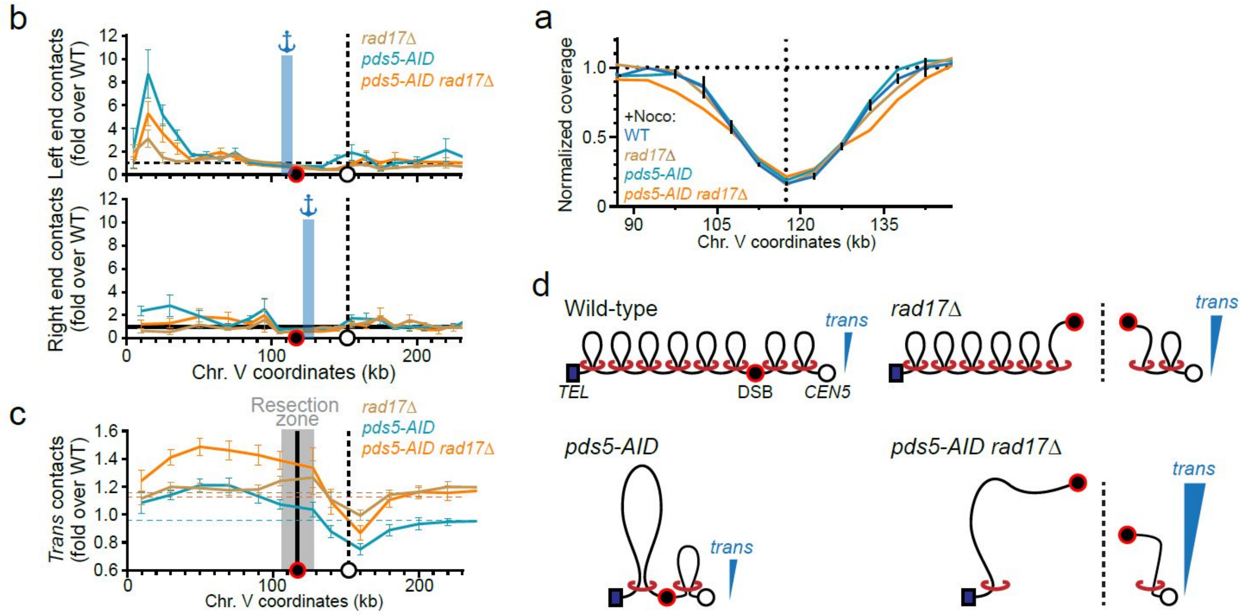


**Figure 9: The 9-1-1 clamp participates in blocking cohesin translocation, inhibiting *trans* contacts by the DSB region.**

**a.** Hi-C coverage binned at 5 kb determined from data in **Fig. 2c, f-g**. Data are mean ±SD, when available.

**b.** 4C-like representation of the fold increase in contacts made by the left (top) and right (bottom) DSB ends in *rad17*Δ, *pds5-AID* and *pds5-AID rad17*Δ mutants relative to a wild-type strain, at 4 hours post-DSB induction on chr V. All strains were treated with nocodazole. White dot: centromere. Red dot: DSB position. Data represent mean ±SEM of ten or twenty 1 kb bins.

**c.** Fold increase in interchromosomal contacts made over the damaged chr. V arm in *rad17*Δ, *pds5-AID* and *pds5-AID rad17*Δ mutants relative to a wild-type strain, 4 hours after DSB induction. Data represent mean ±SEM of twenty 1 kb bins. Horizontal dotted lines represent the genome average of *trans* contacts in each mutant relative to wild-type.

**d.** Model: in wild-type cells, *trans* contacts are limited both by short loop length and 9-1-1-mediated cohesin block at the DSB region. Loss of cohesin block in a *rad17Δ* mutant increases *trans* contacts, limited by overall loop arm length. Increasing loop arm length in a *pds5-AID rad17Δ* mutant further increases the propensity to engage in *trans* contacts.

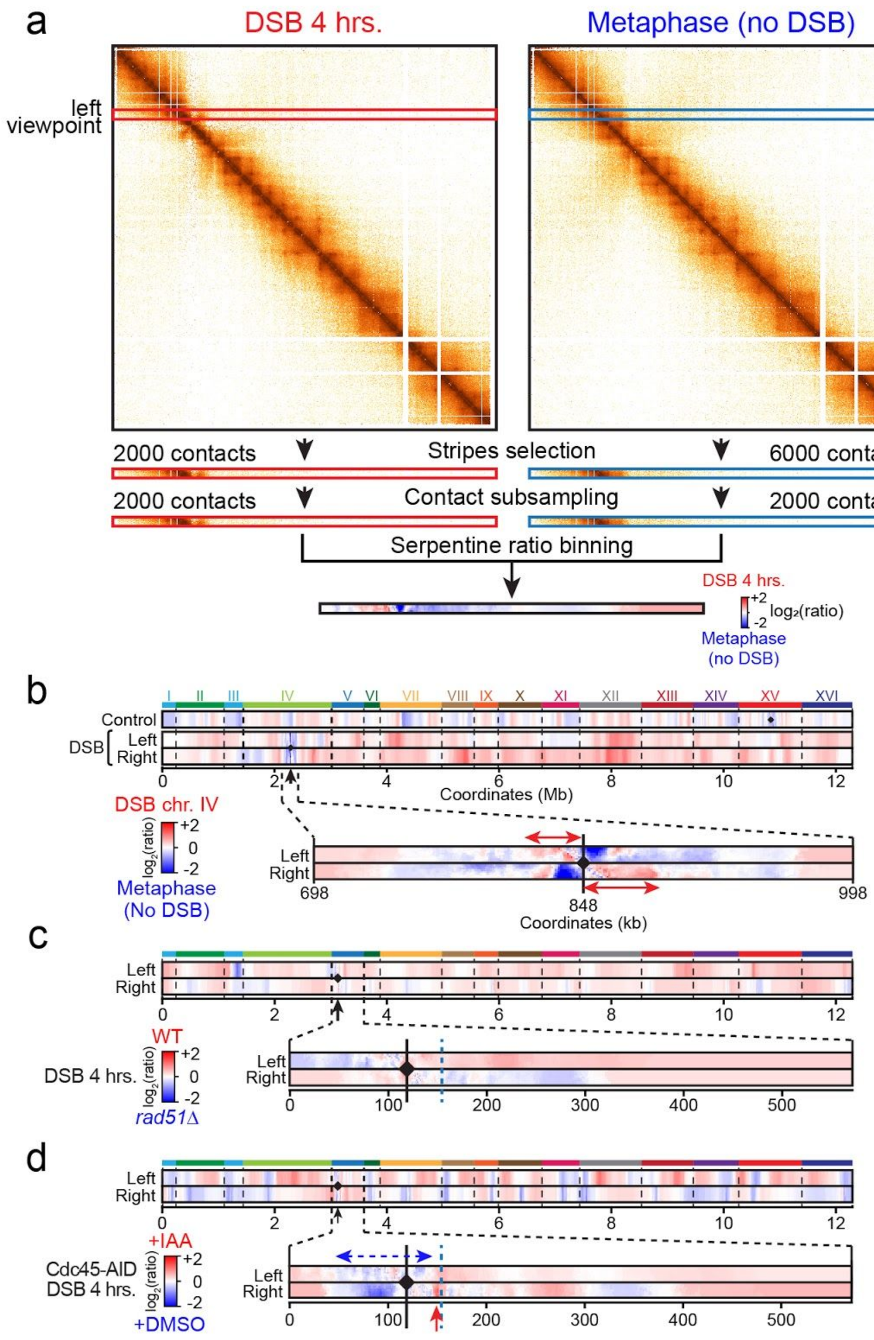


**Figure 10: Study of homology search with Hi-C contact data.**

**a.** Rationale of the differential 4C-like ratio plots in **Fig. 3**. Example of the differential intra-chromosomal contacts made by the left DSB end 4 hrs post-DSB induction (left) over undamaged metaphase-arrested cells (right), from matrices binned at 1 kb. "Stripes" of 10 bins are selected in both matrices and each bin subsampled to match the lowest number of contacts of either matrix. The ratio between both stripes is performed by Serpentine[19], which locally adjusts the binning to the coverage (see **Methods**).

**b.** 4C-like ratio plots of the left and right DSB end on chr IV and a control viewpoint (chr XV) 4 hours post-DSB induction over undamaged metaphase-arrested cells. The genome-wide (top) and intra-chromosomal (bottom) contact profiles are shown. Black arrow: DSB position. Black diamond: location of the control viewpoint.

**c.** 4C-like ratio maps of contact frequencies of the left and right DSB regions 4 hours post-DSB induction in WT over *rad51*Δ cells, genome-wide (top) or within chr. V (bottom).

**d.** Same as (c) but comparing Cdc45-depleted (Cdc45-AID +IAA) over control cells (Cdc45-AID +DMSO) 4 hrs post-DSB induction.

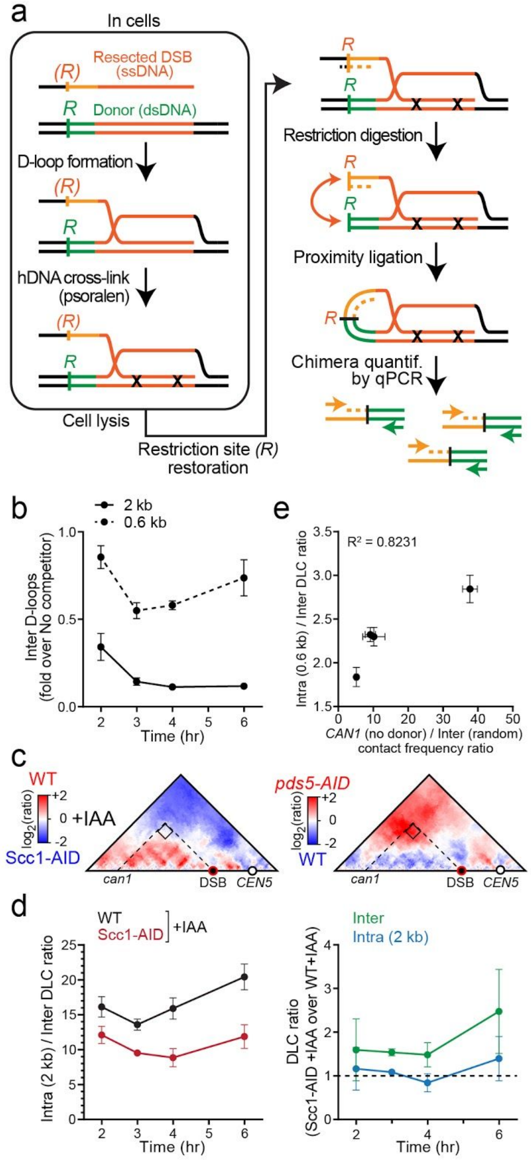


**Figure 11:** **Cohesin promotes homology identification in *cis* and inhibits it in *trans* in multiple ways**

a. Rationale of the DLC assay[12,15]. Following D-loop stabilization by *in vivo* hDNA interstrand crosslink with psoralen, extracted DNA is subjected to endonucleolytic fragmentation with *Eco*RI. Ligation in diluted conditions promotes formation of a chimeric DNA fragment between the physically associated unique sequences flanking the hDNA crosslinked region in the broken and donor molecules. Quantification of this chimera by quantitative PCR is a readout of the amount of D-loops formed at this donor site in the cell population. *R* and *(R)* denote cleavable and uncleavable (*i.e.* in the ssDNA form) *Eco*RI restriction sites, respectively. Restoration of the *Eco*RI site on the resected broken molecule is achieved by providing a complementary oligonucleotide prior to restriction digestion.

b. Inhibition of D-loop formation at the inter-chromosomal donor upon presence of a 0.6 kb- and 2-kb long intra-chromosomal competitor. Data show mean ±SEM of 3 biological replicates.
c. Left panel: Ratio map of wild-type over Scc1-depleted (Scc1-AID+IAA) cells 4 hours post-DSB induction of the left arm of chr V. Right panel: Ratio map of a *pds5-AID* mutant over a wild-type strain 4 hours post-DSB induction of the left arm of chr V. Intersection of dotted lines corresponds to differential contacts between the DSB region and the site used for introduction of the *intra* donor. No homology is present in the strains used to generate these maps.
d. Left panel: ratio of *intra* over *inter* D-loop levels in cells depleted (Scc1-AID + IAA) or not (WT + IAA) for cohesin. Right panel : Fold change of *intra* and *inter* D-loop levels upon Scc1 depletion. The *intra* donor is 2 kb-long. Data show mean ±SEM of three biological replicates.
e. *Intra* over *inter* donor preference as a function of Hi-C contact frequency. DLC and contact data are at 4 hours. Data show mean ±SEM.

**Table S1: Relevant genotype of the haploid Saccharomyces cerevisiae strains used in this study.**

| Strain | Relevant genotype | Source |
|---|---|---|
| APY143 | *MAT***α**-inc, *ura3::loxP, lys2::URA3, trp1::GAL-HO-hphMX, his3-11,15, leu2-3,112, can1-100, ade2-1, RAD5* | This study |
| APY295 | *MAT***α**-inc, *ura3::LY-HOcs, lys2::LY, trp1::GAL-HO-hphMX, his3-11,15, can1-100, leu2-3,112, ade2-1, RAD5* | Piazza et al. 2019 |
| APY331 | *MAT***a**-inc, *ura3::LY-HOcs, lys2::URA3, trp1::GAL-HO-hphMX, his3-11,15, can1-100, leu2-3,112, ade2-1, RAD5* | This study |
| APY332 | *MAT***α**-inc, *ura3::LY-HOcs, lys2::URA3, trp1::GAL-HO-hphMX, his3-11,15, can1-100, leu2-3,112, ade2-1, RAD5* | This study |
| APY428 | *MAT***a**-inc, *ura3::LY-HOcs, lys2::LY, trp1::GAL-HO-hphMX, can1-100, ade2-1, leu2-3,112, his3-11,15, RAD5, sgs1::HIS3* | This study |
| APY429 | *MAT***a**-inc, *ura3::LY-HOcs, LYS2, trp1::GAL-HO-hphMX, can1-100, ade2-1, leu2-3,112, his3-11,15, RAD5, exo1::KMX* | This study |
| APY536 | *MAT***a**-inc, *ura3::LY-HOcs, lys2::LY, trp1::GAL-HO-hphMX, can1-100, ade2-1, leu2-3,112, his3-11,15, RAD5, exo1-D173A::TRP1* | This study |
| APY430 | *MAT***a**-inc, *ura3::LY-HOcs, LYS2, trp1::GAL-HO-hphMX, can1-100, ade2-1, leu2-3,112, his3-11,15, RAD5, exo1::KMX, sgs1::HIS3* | This study |
| APY541 | *MAT***a**-inc, *ura3::LY-HOcs, LYS2, trp1::GAL-HO-hphMX, can1-100, ade2-1, leu2-3,112, his3-11,15, RAD5, mre11::KMX* | This study |
| APY542 | *MAT***a**-inc, *ura3::LY-HOcs, LYS2, trp1::GAL-HO-hphMX, can1-100, ade2-1, leu2-3,112, his3-11,15, RAD5, rad17::KMX* | This study |
| APY525 | *MAT***α**-inc, *ura3::LY-HOcs, lys2::LY, trp1::GAL-HO-hphMX, his3-11,15, can1-100, leu2-3,112, ade2-1, RAD5, rad51::kanMX* | This study |
| APY534 | *MAT***α**-inc, *ura3::LY-HOcs, lys2::LY, trp1::GAL-HO-hphMX, his3-11,15, can1-100, leu2-3,112, ade2-1, RAD5, rad52::kanMX* | This study |
| APY526 | *MAT***a**-inc, *ura3::loxP chrIV-845464::LY-Hocs, lys2::URA3 trp1::GAL-HO-hphMX his3-11,15, leu2-3,112, can1-100, ade2-1, RAD5.* | This study |
| APY531 | *MAT***α**-inc/**a**-inc, *ura3::LY-Hocs/ura3::loxP, lys2::LY/lys2::URA3, trp1::GAL-HO-hphMX/-, his3-11,15/-, can1-100/-, leu2-3,112/-, ade2-1/-, RAD5/-* | This study |
| APY466 | *MAT***α**-inc, *ura3::LY-Hocs, lys2::LY, trp1::GAL-HO-hphMX, can1-100, his3::pADH1-OsTIR1-9Myc::HIS3, ade2-1, leu2-3,112, RAD5* | This study |
| APY467 | *MAT***α**-inc, *ura3::LY-Hocs, lys2::LY, trp1::GAL-HO-hphMX, can1-100, his3::pADH1-OsTIR1-9Myc::HIS3, ade2-1, leu2-3,112, RAD5, scc1-V5-Pk3-AID::kanMX* | This study |
| APY468 | *MAT***α**-inc, *ura3::LY-Hocs, lys2::LY, trp1::GAL-HO-hphMX, can1-100, his3::pADH1-OsTIR1-9Myc::HIS3, ade2-1, leu2-3,112, RAD5, pds5-V5-Pk3-AID::kanMX* | This study |
| APY513 | *MAT***a**-inc, *ura3::LY-Hocs, lys2::LY, trp1::GAL-HO-hphMX, can1-100, his3::pADH1-OsTIR1-9Myc::HIS3, ade2-1, leu2-3,112, RAD5, CDC45-FlagX5-mini-AID::kanMX* | This study |
| APY474 | *MAT***α**-inc, *ura3::LY-HOcs, lys2::LY, trp1::GAL-HO-hphMX, his3-11,15, can1::HIS3-L (0.6kb competitor), leu2-3,112, ade2-1, RAD5* | This study |
| APY296 | *MAT***α**-inc, *ura3::LY-HOcs, lys2::LY, trp1::GAL-HO-hphMX, his3-11,15, can1::HIS3-LY (2kb competitor), leu2-3,112, ade2-1, RAD5* | This study |
| APY473 | *MAT***α**-inc, *ura3::LY-HOcs, lys2::LY, trp1::GAL-HO-hphMX, his3::pADH1-OsTIR1-9Myc::HIS3, can1::HIS3-L (0.6kb competitor), leu2-3,112, ade2-1, RAD5, scc1-V5-Pk3-AID::kanMX* | This study |

| | | |
|---|---|---|
| APY494 | *MAT***α**-inc, *ura3::LY-HOcs, lys2::LY, trp1::GAL-HO-hphMX, his3-11,15, can1::HIS3-L (0.6kb competitor), leu2-3,112, ade2-1, RAD5, pds5-V5-Pk3-AID::kanMX* | This study |
| APY537 | *MAT***a**-inc, *ura3::loxP, lys2::URA3, trp1::GAL-HO-hphMX, his3-11,15, leu2-3,112, can1-100, ade2-1, RAD5, pMET-CDC20::TRP1* | This study |
| APY266 | *MAT***a**-inc, *ura3::LY-HOcs, lys2::LY, trp1::GAL-HO-hphMX, his3-11,15, can1-100, leu2-3,112, ade2-1, RAD5* | This study |
| APY569 | *MAT***a**-inc, *ura3::LY-HOcs, lys2::LY, trp1::GAL-HO-hphMX, his3-11,15, can1-100, leu2-3,112, ade2-1, RAD5, mec3::kanMX* | This study |
| APY597 | *MAT***α**-inc, *ura3::LY-Hocs, lys2::LY, trp1::GAL-HO-hphMX, can1-100, his3::pADH1-OsTIR1-9Myc::HIS3, ade2-1, leu2-3,112, RAD5, PDS5-PK3-AID::KanMX4, rad17::TRP* | This study |
| APY573 | *MAT***a**-inc, *ura3::LY-Hocs, lys2::LY, can1-100, trp1::GAL-HO-hphMX, RAD5,Scc1-PK9::KanMX* | This study |
| APY570 | *alpha-inc ura3::LY-Hocs, lys2::LY, trp1::GAL-HO-hphMX, ade2-1, leu2-3,112, RAD5, can1::HIS3-LY, his3::pADH1-OsTIR1-9Myc::HIS3* | This study |
| APY572 | *MAT***a**-inc, *ura3::LY-HOcs, trp1::GAL-HO-hphMX, leu2-3,112, can1::HIS3-LY, lys2::LY, ade2-1,RAD5, his3::pADH1-OsTIR1-9Myc::HIS3, SCC1-PK1-AID-KanMX* | This study |
| APY540 | *MAT***a**-inc, *ura3::LY-Hocs, lys2::LY, trp1::GAL-HO-hphMX, can1-100, his3::pADH1-OsTIR1-9Myc::HIS3, ade2-1, leu2-3,112, RAD5* | This study |

**Table S2: Primers used in this study.**
**Quantitative PCR primers**

| **Name** | **Sequence (5'-3')** | **Purpose** |
|---|---|---|
| APO79 | AGACAGAATTGGCAAAGATCC | Reference locus (*ARG4*, Ch.VIII); used as a dsDNA loading control. |
| APO80 | GGCCAATTAGTTCACCAAGACG | |
| APO315 | ACTTCGAATTTCGGCACTTC | Quantify the intra-molecular ligation efficiency of a 1904 bp *Eco*RI fragment at the *DAP2* locus. |
| APO316 | CGATGAAACGTTAAGTGACCAC | |
| APO256 | GTTTCAGCTTTCCGCAACAG | Measure dsDNA integrity at the HOcs. DSB induction causes a decrease of signal. |
| APO54 | GGCGAGGTATTGGATAGTTCC | |
| APO55 | AGGAGCACAGACTTAGATTGG | Quantify the *Eco*RI cutting efficiency on the broken molecule (resected) |
| APO139 | AGAGCGGTCAGTAGCAATCC | |
| APO596 | CTTTAACCGGACGCTCGA | Estimate the fraction of resected DNA molecules. Since psoralen does not crosslink ssDNA, the level of amplification relative to the dsDNA (*ARG4*) donor gives an accurate measurement of ssDNA in the cell population. |
| APO597 | TTGAGTTTATTGCTGCCGTC | |
| APO139 | AGAGCGGTCAGTAGCAATCC | Quantify the DLC chimera formed between the upstream unique sequence of the invading molecule with the upstream unique sequence of the donor molecule at the intra-chromosomal (*CAN1*) donor. |
| APO817 | GACGTACAAAGTTCCACTGGC | |

| | | |
|---|---|---|
| APO139 | AGAGCGGTCAGTAGCAATCC | Quantify the DLC chimera formed between the upstream unique sequence of the invading molecule with the upstream unique sequence of the donor molecule at the inter-chromosomal (*LYS2*) donor. |
| APO319 | CACACGCGAAAAACCGCC | |

**Hybridization oligonucleotide**

| **Name** | **Sequence (5'-3')** | **Purpose** |
|---|---|---|
| APO563 (formerly olWDH1770) | CGAAATCATCTTCGGTTAAATCCAAAACGGCAGAAGCCTGAATGAAACATATGAACCAATTGGAGGACGTCAATGAATTCTGGGGATCCATTGCATTTTT | Hybridize at the *Eco*RI site upstream of the PhiX genome fragmenton the resected broken molecule (Ch.V). Heterologous to the DSB construct on its last 5 nucleotides. |
